# Integration of IL-2 and IL-4 Signals Coordinates Divergent Regulatory T cell Responses and Drives Therapeutic Efficacy

**DOI:** 10.1101/2020.04.14.040485

**Authors:** Julie Y. Zhou, Carlos A. Alvarez, Brian A. Cobb

**Affiliations:** Department of Pathology, Case Western Reserve University School of Medicine, Cleveland OH 44106

**Author notes:** Correspondence: Brian A. Cobb, PhD, Professor of Pathology.

**Keywords:** cytokine, T cell, STAT5, STAT6, cytokine receptors, interleukin-2, interleukin-4, interleukin-10, asthma, experimental autoimmune encephalomyelitis

## Abstract

Cells exist within complex milieus of communicating factors, such as cytokines, that combine to generate context-specific responses, yet nearly all knowledge about the function of each cytokine and the signaling propagated downstream of their recognition is based upon the response to individual cytokines. Here, we found that regulatory T cells (Tregs) integrate concurrent signaling initiated by IL-2 and IL-4 to generate a response divergent from the sum of the two pathways in isolation. IL-4 stimulation of STAT6 phosphorylation was blocked by IL-2, while IL-2 and IL-4 synergized to enhance STAT5 phosphorylation, IL-10 production, and the selective proliferation of IL-10-producing Tregs, leading to increased inhibition of conventional T cell activation and the reversal of asthma and multiple sclerosis in mice. These data define a mechanism of combinatorial cytokine signaling and lay the foundation upon which to better understand the origins of cytokine pleiotropy while informing improved the clinical use of cytokines.

**Impact Statement:** Simultaneous cytokine signaling results in unexpected transcription factor changes that fuel a cellular response divergent from the sum of each cytokine alone.

## Introduction

IL-2 and IL-4 are both pleiotropic cytokines and members of a family in which γC (CD132) is a shared receptor subunit. These and many other cytokines have undergone intensive study because of their ubiquitous and essential roles in health and disease (*1*), yet their functions can be highly divergent and context dependent.

IL-2 is critical for launching an effector/memory T cell response as a part of normal immune protection. From a pro-inflammatory perspective, IL-2 can drive the proliferation of activated CD4+ and CD8+ effector T cells (*2, 3*), enhance natural killer cell cytotoxicity (*4, 5*), and augment B cell proliferation and antibody secretion (*6*). However, IL-2 is also potently anti-inflammatory, driving Fas-mediated activation-induced cell death (*7*) while promoting Treg survival (*8*), with its loss resulting in severe autoimmunity and inflammation (*9*). Its pleiotropy can be partly explained by the strategic and context specific expression of its receptors, with IL-2 receptor α (IL-2Rα; CD25) being a common marker for activated conventional T cells (Tconv) as well as an established constitutive marker for Tregs (*10, 11*). CD25 is part of the high affinity receptor complex with IL-2 receptor β (IL-2Rβ; CD122) and γC, although it can function as a low-affinity receptor in isolation (*12*). IL-2 activity is dependent upon Jak/STAT signaling, with STAT3 and STAT5 being the primary mediators of its transcriptional regulation of target genes (*13*).

IL-4 function has perhaps an even broader impact and has two receptor forms. The type 1 IL-4 receptor is comprised of IL-4 receptor α (IL-4Rα; CD124) and γC, whereas the type II receptor contains IL-4Rα and IL-13 receptor α1 (IL-13Rα; CD213A1)(14). Like many cytokines, signaling is mediated by the Jak/STAT pathway, with STAT6 being a main transcriptional mediator (*15*) (*16*). IL-4 is historically associated with allergy through its ability to stimulate class switching to IgE and expression of the IgE receptor (*17*), although it is also critical for parasite clearance (*18*) and differentiation of naïve T cells into a type 2 helper (Th2) phenotype (*19*), all of which is considered pro-inflammatory. Yet, IL-4 is also a key differentiating factor for wound healing macrophages (*20*), which is essential for tissue repair, and is known to oppose IL-17-mediated inflammatory diseases like psoriasis and experimental autoimmune encephalomyelitis (EAE) (*21, 22*). In fact, IL-4 has been shown to elicit IL-10 in both macrophages and T cells (*23, 24*), helping to explain how IL-4 is also potently anti-inflammatory.

Although much is known about the signaling initiated by these and most other cytokines, understanding of how their signals integrate in complex environments remains limited. For many cytokines, the JAK/STAT cascades underlie their transcriptional impact (*25*), although a small number of JAK and STAT proteins belies the divergent nature of the responses. STAT homo- and hetero-multimerization is one mechanism that exponentially expands the available transcription outcomes (*26*). There is also evidence that cytokines can lead to the transcriptional enhancement or diminution of the signaling machinery for other cytokines (*27*). For example, IL-2 stimulation drives the expression of IL-4Rα, thereby enhancing subsequent responses to IL-4 through serial activation of STAT5 and STAT6, respectively, ultimately leading to Th2 T cell differentiation ((*28*)). This is additive signaling, whereby the canonical pathways act in concert to drive T cell differentiation, and follows our knowledge of each pathway in isolation. These mechanisms partially explain how a small number of STAT molecules can drive diverse cellular outputs, but do not account for the possibility of simultaneous stimulation that may occur in the complex array of cytokines that exist in any biological environment.

Here, we have discovered that the combination of IL-2 and IL-4 stimulation in Tregs culminates in a synergistic enhancement of Treg function and proliferation *in vitro* and *in vivo.* This effect is driven by a uniquely integrated response requiring simultaneous rather than serial stimulation. These findings reveal a novel mechanism underlying cytokine signaling which can drive unexpected outcomes that are differential to the simple combination of pathways, thereby laying the groundwork for a more complete picture of cytokine pleiotropy and potentially leading to enhancements to the use of cytokines in the clinical setting.

## Results

### IL-2 and IL-4 synergistically promote IL-10 production by Tregs

Our previous findings demonstrated that IL-4 impacts IL-10 production in FoxP3+ Tregs (*29*), but the nature of the functional interaction between IL-4 and IL-2, a key survival factor for Tregs *in vitro* and *in vivo,* remained obscure. Thus, we created a novel dual reporter mouse by crossing FoxP3^RFP^ (*30*) and IL-10^GFP^ (*31*) mice, thereby enabling live sorting of FoxP3+ cells and analysis of IL-10 production on a per-cell basis. CD4+FoxP3+ Tregs isolated from the spleens of naïve dual reporter mice (Figure 1 – figure supplement 1A-B) by magnetic bead and sterile fluorescence-activated cell sorting (FACS) were cultured with T cell receptor (TCR) activation using αCD3ε antibody and all combinations of IL-2 and IL-4 for 3 days. We found that Tregs cultured with combinatorial cytokine stimulation resulted in synergistically higher numbers of IL-10 expressing cells (Figures 1A–1C) and IL-10 secretion (Figure 1D) compared to single cytokine stimulation. However, analysis of IL-10^+^ cells revealed that IL-10 expression was no different than IL-2-stimulation in isolation (Figure 1E), suggesting that the cytokines in combination do not elicit a synergistic increase in IL-10 production on a per-cell basis. The sex-independent (Figure 1F) and TCR-stimulation dependent synergy (Figures 1A–1D) was present in FoxP3+ Tregs but not FoxP3^-^ Tconv (Figure 1A), and no loss of FoxP3 expression was observed (Figure 1G), suggesting that the machinery required for this effect was unique to FoxP3^+^ Tregs.

**Figure 1.**
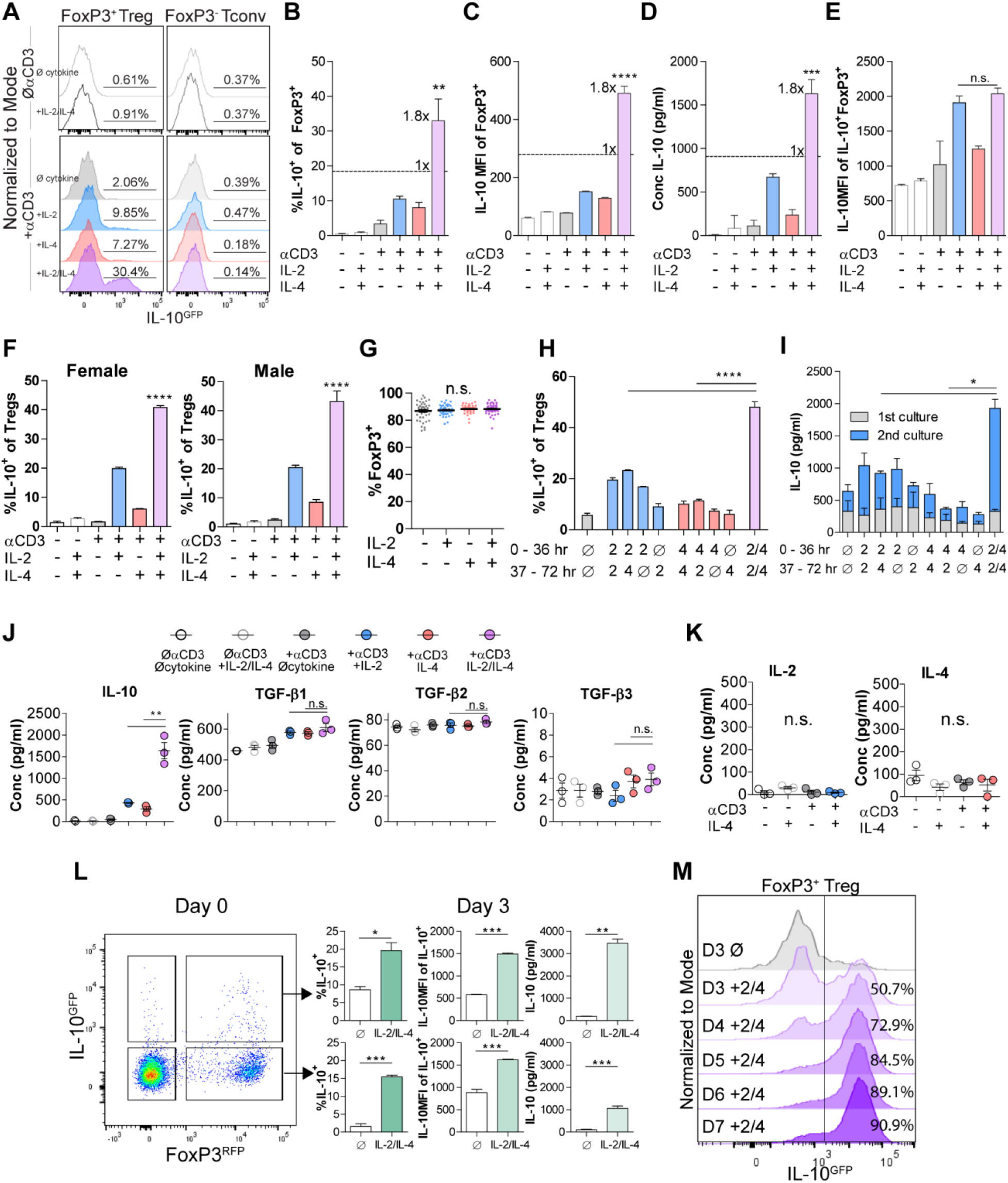
IL-2 and IL-4 synergistically promote IL-10 production by Tregs. (A-C) IL-10 expression of Tregs purified from FoxP3^RFP^/IL-10^GFP^ dual reporter mice (See also Figure 1 – figure supplement 1) cultured for 3 days with the designated stimulants as analyzed by flow cytometry. For all panels, N≥3 for all bar graphs and histograms are representative. (D) IL-10 production of Tregs cultured with the designated stimulation as quantified by ELISA of the culture supernatants. N=3. (E) IL-10 expression of purified IL-10^+^ Tregs cultured with the designated stimulation as quantified by flow cytometry. N=3 (F) Female and male responses to combinatorial cytokine stimulation after 3 days, as measured by flow cytometry for IL-10 expression. N=3. (G) FoxP3 expression by purified Tregs stimulated in culture for 3 days with the designated conditions, as analyzed by flow cytometry. N≥27. (H) IL-10 expression of purified Tregs stimulated for 36 hours in culture, washed, then subsequently stimulated for another 36 hours in culture with the indicated conditions, as analyzed by flow cytometry. All samples received αCD3ε activation. (See also Figure 1 – figure supplement 3A). N=3 (I) IL-10 production by purified Tregs stimulated for 36 hours in culture, washed, then subsequently stimulated for another 36 hours in culture with the indicated conditions, as analyzed by ELISA of the culture supernatants. All samples received αCD3ε activation. (See also Figure 1 – figure supplement 3B). N=3 (J) Cytokine production following 3 days of Treg culture as quantified by multianalyte Luminex of the culture supernatants (See also Figure 1 – figure supplement 2). N=3 (K) IL-2 and IL-4 production by Tregs following 3 days of stimulation with the designated conditions, as quantified by ELISA of the culture supernatants. N=3 (L) IL-10 production of purified IL-10^+^ or IL-10^-^ Tregs following 3 days of culture with αCD3ε and combined IL-2/IL-4, as analyzed by flow cytometry. (See also Figure 1 – figure supplement 3C). N=3 (M) IL-10 production of purified Tregs cultured with αCD3ε and combined IL-2/IL-4 for 3 to 7 days as analyzed by flow cytometry (See also Figure 1 – figure supplement 3D). Histograms are representative of 3 independent experiments. Mean ± SEM are indicated. *p<0.05, **p<0.01, ***p<0.001, ****p<0.0001

In order to determine whether the cytokines must be present at the same time, we isolated Tregs and stimulated them with cytokines in series over a 3-day culture and compared the response to simultaneous stimulation. We discovered that both IL-2 and IL-4 must be present in culture together to manifest the synergistic IL-10 response (Figures 1H-I and Figure 1 – figure supplement 3A-B), implicating the integration of IL-2 and IL-4-mediated intracellular signaling as a key underlying mechanism.

To ascertain whether IL-2/IL-4 also affects the ability of Tregs to produce other cytokines and chemokines, Treg supernatants after 3 days of culture with or without TCR stimulation and all cytokine combinations were analyzed by multianalyte Luminex. Remarkably, no cytokine or chemokine other than IL-10 was synergistically and robustly increased (Figure 1 – figure supplement 2) by combined IL-2 and IL-4, including the Treg-associated TGF-β family members (*32*)(Figure 1J). Moreover, neither IL-2 nor IL-4 induced autocrine production of the reciprocal cytokine (Figure 1K).

To differentiate whether the combined cytokines induced or maintained IL-10 expression in Tregs, we sorted splenocytes from the dual reporter mice into FoxP3+IL-10^-^ and FoxP3+IL-10^+^ populations, then cultured them in all combinations. We observed robust induction of IL-10 expression in FoxP3+IL-10^-^ cells and maintenance of IL-10 expression in FoxP3+IL-10^+^ cells, with no detectable change in FoxP3 (Figures 1L and Figure 1 – figure supplement 3C). To assess this effect beyond 3 days, we cultured Tregs and analyzed them on days 3-7 and found a profound daily increase in the overall percentage of IL-10^+^ Tregs, ultimately leading to nearly every Treg converting to IL-10^+^ by day 7 (Figures 1M and Figure 1 – figure supplement 3D).

### IL-2/IL-4 enhances Treg proliferation and selectively drives the expansion of IL-10^+^ Tregs

The lack of a major difference in per-cell IL-10 expression after exposure to IL-2 and IL-4 together (Figure 1E) coupled with the conversion of nearly all Tregs into IL-10^+^ cells (Figure 1M) and synergistic IL-10 release (Figure 1D) suggested that the cells were proliferating in response to the combined cytokines. Dual reporter Tregs were isolated and stained with CellTrace Violet and stimulated with all cytokine combinations for 3 days. Tregs cultured with the combined cytokines were more proliferative, as measured by the number and magnitude of cell division peaks (Figure 2A), proliferation index, and division index (Figure 2B). The division index revealed that more Tregs stimulated by the combined cytokines divide, and the proliferation index indicated that each dividing Treg undergoes more divisions than when stimulated with or without the single cytokines.

**Figure 2.**
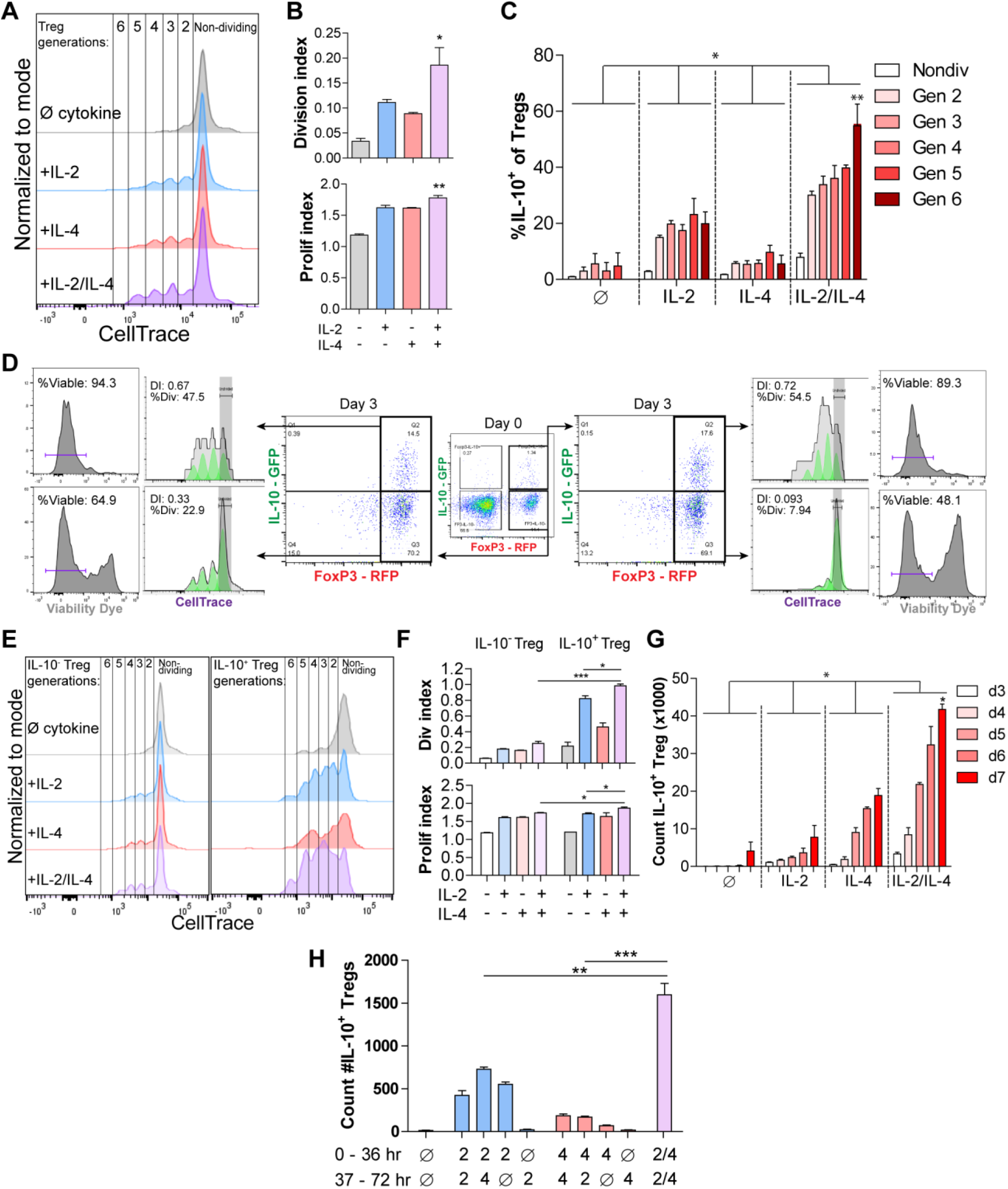
IL-2/IL-4 enhances Treg proliferation and selectively drives the expansion of IL-10^+^ Tregs. (A-B) Treg proliferation as measured by CellTrace signal following 3 days of culture with αCDЗε and all combinations of IL-2 and IL-4, with Division and Proliferation Indices indicated. N=3 for all bar graphs and histograms are representative. (C) IL-10 expression of purified Tregs cultured for 3 days with αCD3ε and the designated cytokines, and gated by CellTrace generation, as measured by flow cytometry (panel A). N=3. The IL-10 expression of all IL-2/IL-4-induced Treg generations are statistically significant (p <0.05) compared to the other cytokine-stimulated conditions. (D) Proliferation and viability of purified IL-10^+^ and IL-10^-^ Tregs cultured for 3 days with αCDЗε and both IL-2 and IL-4, as flow cytometry analysis of CellTrace and Sytox Red signal. The histograms are representative. N=3 (E-F) Proliferation of purified of Tregs cultured for 3 days with αCDЗε and all cytokine combinations, gated on IL-10^+/-^ expression, as analyzed by flow cytometry. N=3 for all bar graphs and histograms are representative. (G) Cell count of IL-10^+^ Tregs following 3-7 days of culture with αCD3ε and all combinations of IL-2 and IL-4, as analyzed by flow cytometry. N=3. The cell counts of all IL-2/IL-4-induced conditions are statistically significant (p <0.05) compared to the other cytokine-stimulated conditions. (H) Enumeration of purified Tregs that underwent stimulation with αCD3ε and all cytokine combinations for 36 hours in culture with the first stimulation conditions, and another 36 hours with the second stimulation conditions after washing. Cells underwent flow cytometry. N=3. For all panels, mean ± SEM are indicated. *p<0.05, **p<0.01, ***p<0.001

Since IL-2 and IL-4 in combination elicited synergistic IL-10 production by Tregs (Figure 1), we determined whether Treg proliferation and viability was correlated with higher IL-10 expression among dividing cells. We found that cytokine supplementation led to each subsequent generation of Tregs expressing more IL-10 than the previous generation (Figure 2C). Furthermore, by staining IL-10^-^ and IL-10^+^ Tregs with CellTrace prior to culturing and Sytox viability dye after culturing revealed that originally IL-10^-^ Tregs that gained IL-10 expression adopted a higher division index than the Tregs which remained IL-10^-^ (Figure 2D), and Tregs that were IL-10^+^ on day 0 and remained IL-10^+^ after 3 days of culture had the highest division index and percentage of the overall population that divided. Lastly, the IL-10^+^ Tregs that lost IL-10 expression had a drastic decrease in proliferation as measured by division index and percentage of cells that underwent division, indicating that IL-10^+^ Tregs only lose IL-10 expression when they become quiescent and non-dividing.

We then compared the division and proliferation indices of IL-10^+^ with IL-10^-^ Tregs, and found that IL-10-expressing Tregs were indeed dramatically more proliferative than Tregs that do not express IL-10 (Figure 2E), and that IL-10^+^ Tregs supplemented with combined IL-2 and IL-4 proliferated the most of all. Moreover, the increased proliferation and division indices in IL-2/IL-4-treated IL-10^+^ Tregs compared to IL-10^-^ Tregs suggests that expression of IL-10 in Tregs, as induced by the combined cytokines, coincides with more cell proliferation (Figure 2F).

Finally, in a time course study where purified Tregs were cultured in the presence of all cytokine combinations for 3-7 days, the combined IL-2 and IL-4 synergistically increased the number of IL-10^+^ Tregs compared to single cytokines, most notably increasing the counts of IL-10^+^ Tregs over 12-fold in 5 days (Figures 2G). As seen with IL-10 induction (Figure 1H), exposing cells to the cytokines in series rather than together failed to duplicate the proliferative response (Figure 2H).

Collectively, these data (Figures 1–2 and Figure 1 – figure supplements 1–3) indicate that not only does the combination of IL-2 and IL-4 lead to strong IL-10 expression and enhanced proliferation among TCR-activated Tregs, but that these events are associated in such a way that leads to the selective propagation of IL-10^+^ Tregs resulting in an exponentially increased Treg response.

### Combined IL-2 and IL-4 increase the suppressive ability of Tregs

The ability of IL-10 to suppress the proliferation and functions of conventional and effector T cells is well known (*33*). Since the cytokines working together elicit extraordinarily robust IL-10 production by Tregs through a combination of IL-10 induction and cellular proliferation, we assessed the *in vitro* suppressive impact on Tconv proliferation and cytokine production. Tregs were stimulated as before for 3 days, washed, counted, and normalized to the number of surviving Tregs, and placed them in coculture with freshly purified and CellTrace-stained Tconv cells harvested from the spleens of dual reporter mice. After 3 days, the cells were analyzed for proliferation and cytokine production. Treg suppression activity was impaired upon IL-4 stimulation, despite the IL-4-mediated increase in IL-10 (Figure 1), while supplementation with either IL-2 or both cytokines substantially increased Tconv inhibition (Figure 3A). Stimulation by combined IL-2 and IL-4 led to increased IL-10 release by Tregs (Figures 3B) and significant suppression of IL-4 and IL-17A production by Tconv cells (Figures 3C), but only a modest increase in suppressive capacity over IL-2 exposure alone.

**Figure 3.**
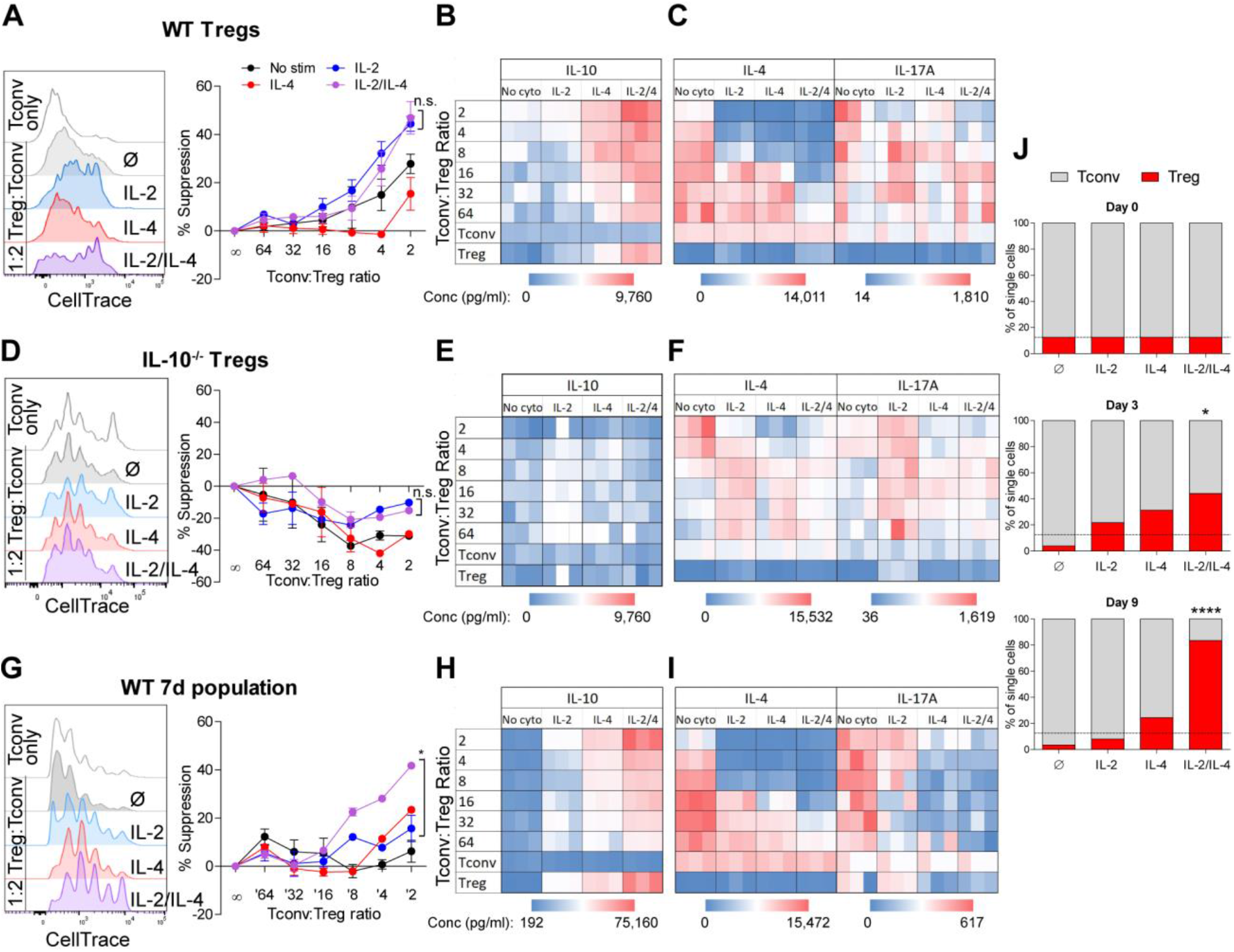
Combined IL-2 and IL-4 increase the suppressive ability of Tregs. (A-C) Proliferation (flow cytometry) and cytokine output (ELISA) of freshly isolated and Celltrace-stained Tconv co-cultured with αCDЗε and the indicated ratios of WT Tregs separately and previously stimulated with αCDЗε and all cytokine conditions for 3 days and normalized for cell number at the time of coculture. N=3 for all graphs and histograms are representative. (D-F) Proliferation (flow cytometry) and cytokine production (ELISA) of freshly isolated Tconv cells cocultured with αCDЗε and the indicated ratios of IL-10^-/-^ Tregs separately and previously stimulated with αCDЗε and all cytokine conditions for 3 days and normalized for cell number at the time of co-culture. N=3 for all graphs and histograms are representative. (G-I) Proliferation (flow cytometry) and cytokine production (ELISA) of freshly isolated Tconv cells cocultured with αCDЗε and the indicated ratios of WT Tregs separately and previously stimulated with αCDЗε and all cytokine conditions for 7 days. Co-culture ratio was based on the number of Tregs prior to cytokine stimulation to incorporate their proliferation. N=3 for all graphs and histograms are representative. (J) Percentage of the overall culture comprising of Tregs and Tconv as determined by flow cytometry over the course of time. The Tregs were purified and stimulated with αCDЗε and all cytokine conditions for 3 days, washed, then placed in co-culture with freshly isolated Tconv cells at a 1:8 Treg:Tconv starting ratio (line). N=3 For all panels, mean ± SEM are indicated. *p<0.05, ****p<0.0001

Next, to assess the role of IL-10 in Tconv inhibition, we repeated the experiment using Tregs isolated from IL-10-deficient (IL-10^-/-^) mice and found that cytokine-stimulated IL-10^-/-^ Tregs failed to suppress Tconv proliferation (Figure 3D), did not express IL-10 (Figures 3E), and failed to reduce Tconv cytokine expression (Figures 3F). This demonstrates that IL-10 is necessary for cytokine-stimulated Tregs to suppress Tconv cells.

These data show that on a per-cell basis, Tregs stimulated with IL-2 alone have similar suppressive activity compared to the IL-2/IL-4 combination, and that this activity is IL-10 dependent. This agrees with the lack of difference in per-cell IL-10 production (Figure 1E), but fails to account for the enhanced and selective proliferation of IL-10^+^ Tregs with combinatorial cytokine exposure. To do this, we cultured Tregs alone with all combinations of cytokines for 7 days (Figures 3A–3C) and washed them after stimulation, but did not re-normalize the cell numbers prior to co-culture with fresh Tconv cells to allow for proliferative differences to be incorporated into the activity assessment. We observed that Tregs cultured without cytokine had lost their suppressive ability, likely due to an extended amount of time without supportive growth signals in culture, and that the Tregs cultured with single cytokines maintained suppressive activity (Figures 3G–3I). However, Tregs stimulated with both cytokines showed significantly greater suppression of Tconv proliferation (Figure 3G), produced higher concentrations of IL-10 (Figures 3H), and suppressed the production of IL-4 and IL-17A by Tconv cells to a greater degree (Figures 3I) compared to single cytokine conditions. Indeed, we found that by day 9 of a 1:8 Treg-Tconv co-culture, the Tregs previously stimulated with both IL-2 and IL-4 comprised over 80% of the resulting culture (Figure 3J), demonstrating that the synergistic impact of these cytokines is manifested through the combination of IL-10 production, suppression of Tconv proliferation and cytokine production, and the selective proliferation of IL-10^+^ Tregs even after removal from recombinant cytokine stimulation.

### Synergistic IL-10 production and proliferation is STAT5-dependent

The intracellular signaling downstream of IL-2 and IL-4 receptors is well known, including STAT6 as characteristic of IL-4 signaling (*34*) and STAT5 being associated with IL-10 production in Tregs downstream of IL-2 (*35*). In order to understand the mechanism by which IL-2 and IL-4 signals are integrated, we measured STAT phosphorylation by flow cytometry following Treg stimulation as before. While STAT3 is associated with both IL-2 and IL-4, we found that STAT3 phosphorylation was not altered as a result of combined cytokine stimulation in TCR-activated Tregs, although a small amount of STAT3 phosphorylation occurred at 60 minutes with combined cytokines without TCR stimulation (Figure 4A). IL-2 induced low levels of STAT6 phosphorylation, while IL-4 induced robust STAT6 phosphorylation in TCR-activated Tregs. Remarkably, the addition of IL-2 to the IL-4 and TCR signals resulted in the reversal of IL-4-dependent STAT6 activation (Figure 4B). Mirroring IL-10 production, we also detected a dramatic and synergistic increase in STAT5 phosphorylation when TCR-activated Tregs were given both cytokines (Figure 4C), suggesting that STAT5 may be responsible for the synergistic IL-10 and proliferative response.

**Figure 4.**
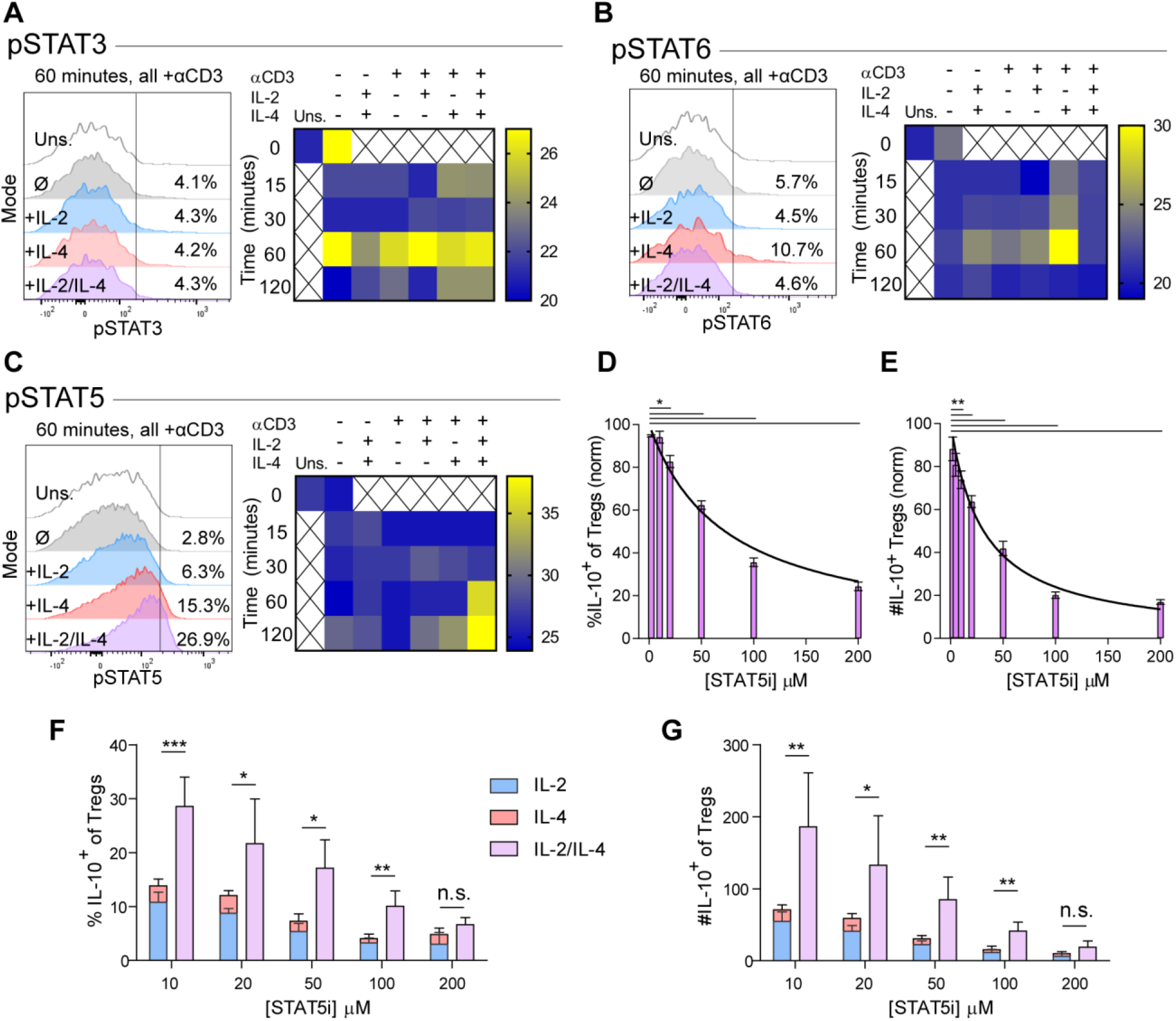
Synergistic IL-10 production and proliferation is STAT5-dependent. (A-C) pSTAT3, pSTAT6, and pSTAT5 expression of purified Tregs stimulated with the indicated culture conditions as analyzed by flow cytometry. The histograms are representative and heatmaps represent N≥3 experiments. (D-E) Treg expression of IL-10 and enumeration of IL-10^+^ Tregs following αCD3ε, combined IL-2/IL-4, and STAT5i supplementation in 3 day culture, as analyzed by flow cytometry (See also Figure 4, figure supplement 1). N=3. (F-G) IL-10 expression in Tregs and IL-10^+^ Tregs counts following αCD3ε, all combinations of IL-2 and IL-4, and STAT5i supplementation in 3 day culture, as analyzed by flow cytometry. N=3. For all panels, mean ± SEM are indicated. *p<0.05, **p<0.01, ***p<0.001

In order to determine whether STAT5 activation is a point of signaling convergence necessary for synergy, we blocked STAT5 activity over a range of increasing inhibitor (STAT5i) doses in stimulated Tregs. Blockade of STAT5 activity inhibited the response by over 80 % in a dose-dependent fashion, as measured by inhibition of IL-10 induction as a percent (Figure 4D) and number (Figure 4E) of IL-10^+^ Tregs, overall Treg viability (Figure 4 – figure supplement 1A), IL-10 expression on a cellular level (Figure 4 – figure supplement 1B), proliferation (Figure 4 – figure supplement 4C-D), and IL-10 release (Figure 4 – figure supplement 1E). As a control, analysis of IFNγ, a cytokine not produced synergistically following combined IL-2 and IL-4 stimulation, demonstrated no inhibition (Figure 4 – figure supplement 1F).

**Figure 5.**
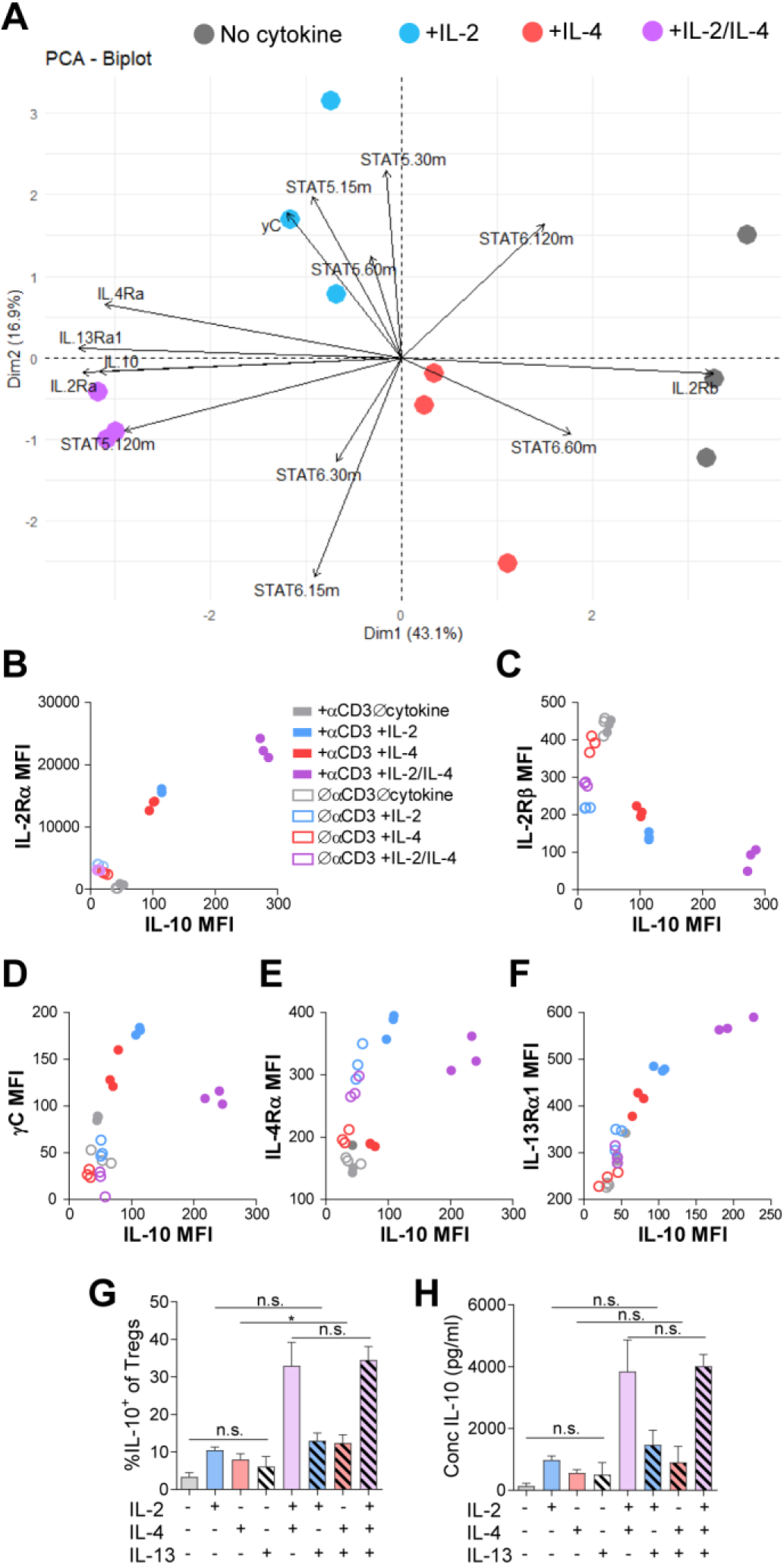
Combinatorial IL-2 and IL-4 signaling promotes the expression of the Type 2 IL-4 receptor. (A) Expression of 14 parameters as shown in an unpaired principle component analysis of Tregs purified and stimulated with αCDЗε and all cytokine conditions for З days unless otherwise noted in the vectors. N=3 (B-F) Surface expression of receptor subunits that comprise the multiple forms of the IL-2R and IL-4 on purified Tregs from FoxP3^RFP^/IL-10^GFP^ dual reporter mice that were stimulated for 3 days with or without αCDЗε and with all cytokine combinations. Expression of receptor subunits is plotted as a function of IL-10 expression. N=3 (G-H) Quantification of IL-10 expression and protein concentration secreted by Tregs that were purified and cultured for 3 days with αCD3ε and all combinations of IL-2, IL-4, and IL-13, as analyzed by flow cytometry and ELISA, respectively. N=3. For all panels, mean ± SEM are indicated. *p<0.05

These data demonstrated that IL-10 production and proliferation downstream of IL-2 and IL-4 was dependent upon STAT5; however, it remained unclear whether cytokine synergy also required STAT5. We again cultured Tregs with the single or combinatorial cytokines and a range of increasing concentrations of STAT5i. Although STAT5i inhibited IL-10 expression and reduced the number of IL-10^+^ Tregs under all conditions (Figure 4F–4G), we discovered that at high STAT5i concentration, IL-2/IL-4 synergy is lost, as illustrated by a lack of difference between the sum of the individual IL-2 and IL-4 responses (Figures 4F–4G; stacked bars) and stimulation with both cytokines simultaneously (Figures 4F–4G; purple bars).

These data demonstrate that the signaling downstream of individual cytokine receptors can be altered by the combinatorial integration of other cytokine-stimulated signals, as is clear for STAT5 and STAT6. Moreover, it is clear that not only are IL-10 production and Treg proliferation paired events, but that STAT5 is necessary to induce the synergistic response characteristic of the IL-2 and IL-4 combination.

### Combinatorial IL-2 and IL-4 signaling promotes the expression of the Type 2 IL-4 receptor

We have found that IL-2 and IL-4 in combination promotes selective proliferation of IL-10^+^ Tregs (Figure 2), which suggests that the receptor expression may change over time to promote the maintenance of IL-10 expression. Thus, we measured cell surface expression of all IL-2 and IL-4 receptor subunits. Tregs were again stimulated, and each subunit was quantified by flow cytometry (Figure 5). Surface expression was first visualized using principle component analysis (PCA), which demonstrates that the receptor expression pattern of TCR-activated Tregs without cytokines, with IL-2 or IL-4 individually, and both cytokines are all divergent and distinct (Figure 5A), further supporting a model in which Tregs possess the ability to integrate multiple cytokine signals in ways distinct from simply adding known pathways studied in isolation. Further analyses of the data revealed a strong positive correlation between IL-10 and IL-2Rα (CD25) expression (Figure 5B), which agrees with both a prior study which reported STAT5-dependent expression of IL-2Rα on Tregs (*36*) and our STAT5 data (Figure 4C). We also discovered a strong negative correlation between IL-10 and IL-2Rβ expression (Figure 5C), and although there is a positive correlation between IL-10 and γC chain in cells receiving individual cytokines, this is significantly reduced when IL-2 and IL-4 are together (Figure 5D). Looking at the IL-4 receptor subunits, the IL-4Rα expression pattern (Figure 5E) mirrors that observed for the γC chain (Figure 5D), with a positive correlation with IL-10 for the individual cytokines, but less so with the combination. Curiously, IL-13Rα1 expression showed a strong positive correlation with IL-10 (Figure 5F), similar to IL-2Rα (Figure 5B), following combined cytokine stimulation. Since the type II IL-4R is used by both IL-4 and IL-13, we tested whether IL-13 could achieve the same synergistic Treg outcome as IL-4 in combination with IL-2. Supplementing Treg cultures with recombinant IL-13 in any combination with IL-2 and IL-4 did not impact IL-10 production (Figures 5G-H), suggesting that the response is specific for IL-4. These data collectively demonstrate that IL-2Rα and the type II IL-4R (IL-4Rα + IL-13Rα) are selectively associated with the synergistic response to the IL-2 and IL-4 combination.

### Combinatorial IL-2 and IL-4 signaling suppresses asthma-like pulmonary morbidity

Given the potent *in vitro* Treg response to the combination of IL-2 and IL-4 and the resulting integration of their signaling pathways, we next sought to determine the *in vivo* response within the context of inflammatory disease. Since this pathway was implicated using murine asthma (*29, 37–39*), we first challenged mice with house dust mite (HDM) over the course of 2 weeks to induce pulmonary inflammation as described previously (*29*). The combinatorial cytokines were administered i.n. at 3 time points prior to HDM challenge as a prophylactic therapy (Figure 6 – figure supplement 1A). We found that mice pretreated with a 1:10 ratio of IL-2:IL-4 were significantly protected from developing airway disease as quantified by leukocyte airway infiltration in bronchial alveolar lavage fluid (BALf)(Figure 6A). Tissue histology by H&E staining further demonstrated a reduction of cellular infiltration around the major airways and vasculature, and decreased bronchial epithelial cell hyperplasia (Figure 6B), while PAS staining revealed reduced mucus production (Figure 6C). Flow cytometry of lung and spleen cells showed that the mice administered combined cytokines accumulated more IL-10^+^ and total Tregs per gram of tissue (Figure 6D). The cytokines were not able to reduce disease morbidity in mice deficient in IL-10 as indicated by BALf cell differentials and lung histology (Figure 6 – figure supplement 1B). Indeed, IL-10 deficient mice appeared to trend towards more severe disease with cytokine administration.

**Figure 6.**
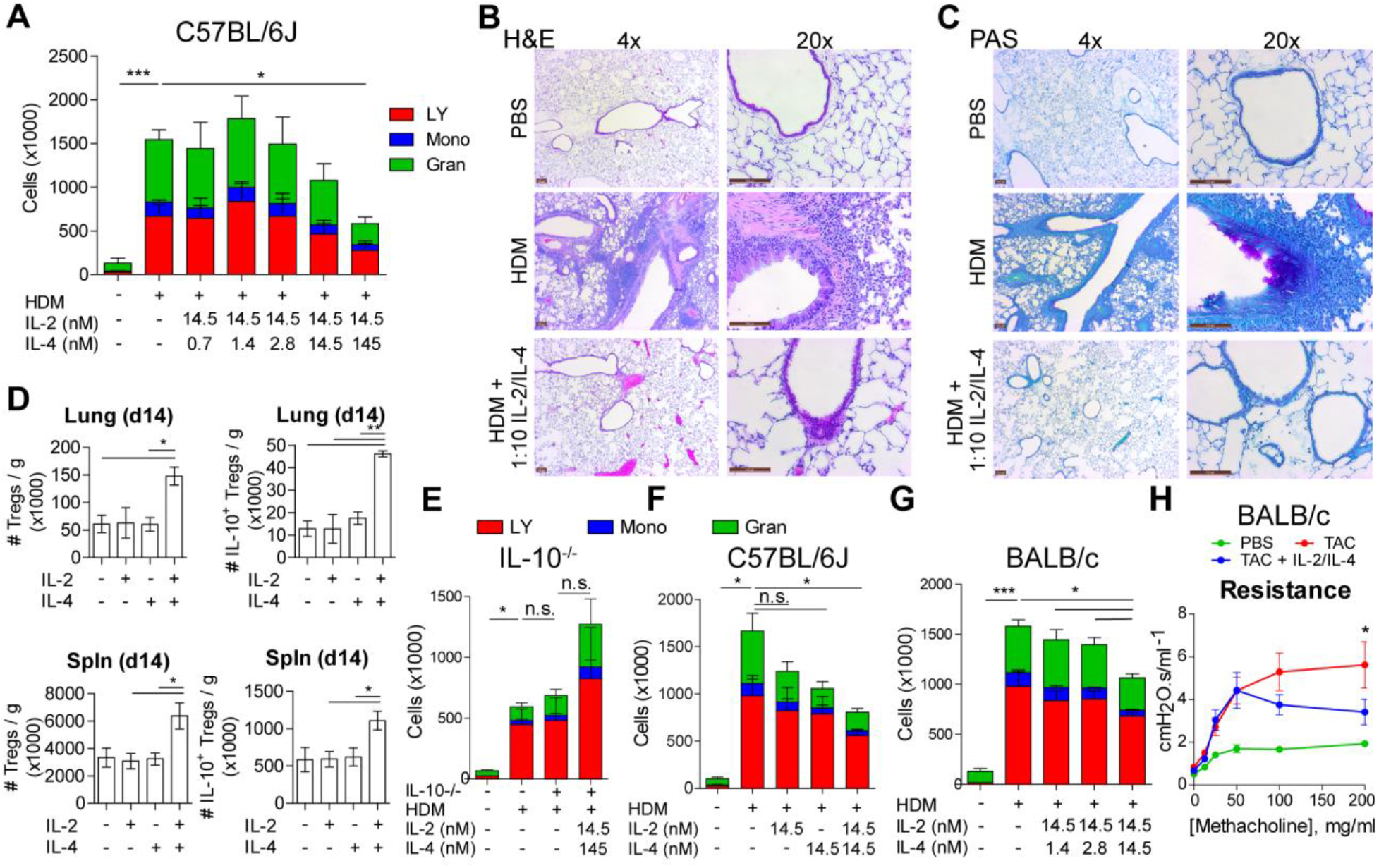
IL-2 and IL-4 in combination suppress the severity of HDM-induced asthma. (A-C) Analysis of HDM-induced airway inflammation in WT C57Bl/6J mice by cell differentials of the Bronchial Alveolar Lavage fluid (BALf) and H&E and PAS staining of FFPE-processed pulmonary tissues. IL-2 and IL-4 were administered i.n. on days 0-5. See also Figure Figure 6 – figure supplement 1A. N=6. Histological images are representative. (D) Quantification of Treg and IL-10^+^ Treg numbers in naïve mice following i.n. administration of combined IL-2 and IL-4 on days 0-5. Mice were harvested at day 14 for flow cytometric analysis. N=6. (E) Assessment of airway inflammation in IL-10^-/-^ C57Bl/6J mice by cell differential analysis following i.n. challenge with HDM and i.n. administration of combined IL-2 and IL-4 in a preventative regiment in the indicated doses (See also Figure 6, figure supplement 1B). N=6. (F-G) Cell differential analysis of BALF quantifying the severity of HDM-induced airway inflammation in mice (C57Bl/6J or BALB/c as indicated) and i.n. administration of cytokine on days 7-11 as indicated (See also Figures 6, figure supplement 1C-E). N=6. (H) Airway resistance in chronic triple antigen-challenged (TAC; HDM/CRA/OVA) BALB/c mice treated or not with i.n. IL-2 and IL-4 starting in week 3 (See also Figure 6, figure supplement 2). Airway resistance was measured by Flexivent as a dose response to methacholine. N≥2. For all panels, mean ± SEM are indicated. *p<0.05, **p<0.01, ***p<0.001

We further investigated the efficacy of the combined cytokines as therapy to ongoing disease rather than as a prophylaxis (Figure 6 – figure supplement 1C). We found that a 1:1 ratio of IL-2/IL-4 was sufficient to suppress disease, as indicated by reduced BALf cell differentials (Figure 6F) and greatly improved lung histology (Figure 6 – figure supplement 1D). In comparison, individual cytokines administered at the same doses failed to significantly suppress infiltration (Figure 6F) or tissue pathology (Figure 6 – figure supplement 1D).

Since asthma is often associated with IL-4 and type 2 immunity, we used the therapeutic treatment regimen (Figure 6 – figure supplement 1C) in the classically Th2-skewed BALB/c mouse background. Remarkably, IL-2 and IL-4 administration in combination significantly suppressed disease when given at the equimolar dose (Figures 6G and Figure 6 – figure supplement 1E), demonstrating that these cytokines do not exacerbate asthma pathology even in a Th2-skewed system.

Finally, asthma in humans often presents as a chronic disease that affects patients for the duration of their life. We therefore tested the efficacy of the cytokines in a chronic model of pulmonary inflammation induced by challenging mice with three alternating antigens (i.e. HDM, cockroach antigen, and ovalbumin) over 10 weeks (Figure 6 – figure supplement 2A). Since this model results in tissue remodeling and hyper-responsiveness to bronchial restricting agents, we examined the lung function of the mice using a Flexivent ventilator, and observed that the mice treated with the combined cytokines had significantly lower lung resistance upon methacholine challenge (Figure 6H). Moreover, lung histology revealed reduced cellular infiltration and epithelial hyperplasia by H&E staining, decreased mucus production (PAS staining), and prevention of long-term tissue remodeling (TriChrome staining) when provided IL-2 and IL-4 combination therapy (Figure 6 – figure supplement 2B).

These data reveal that the combination of IL-2 and IL-4 serves as a robust preventative and therapeutic treatment for multiple models of murine asthma on both Th1 and Th2-skewed backgrounds.

### Combinatorial IL-2 and IL-4 signaling suppresses EAE morbidity

Since the asthma models allow for direct administration of cytokine to the site of inflammation via inhalation, we sought to determine whether systemic administration of both cytokines would also be beneficial in the IL-17-driven model EAE. EAE was induced in dual reporter mice by sensitization with myelin oligodendrocyte glycoprotein (MOG)_35-55_ peptide and adjuvant. When the combined cytokines were injected i.v. into mice challenged with MOG, all three delivery schedules (Figure 7 – figure supplement 1A), including preventative (i.e. prophylactic), concomitant (i.e. at the first sign of disease), and therapeutic (i.e. approaching peak disease), generated significant reductions in clinical score (Figures 7A and Figure 7 – figure supplement 1B). Histological analyses indicated that mice treated with the combined cytokines had reduced cellular infiltration (Figures 7B and Figure 7 – figure supplement 1C; H&E) and demyelination (Figures 7B and Figure 7 – figure supplement 1C; Luxol Fast Blue) of the spinal cord. Flow cytometry further revealed that the cytokines led to significantly reduced MOG-specific Tconv cells (Figure 7C) while ELISA showed decreased IL-17 within neuronal tissues (Figure 7D).

**Figure 7.**
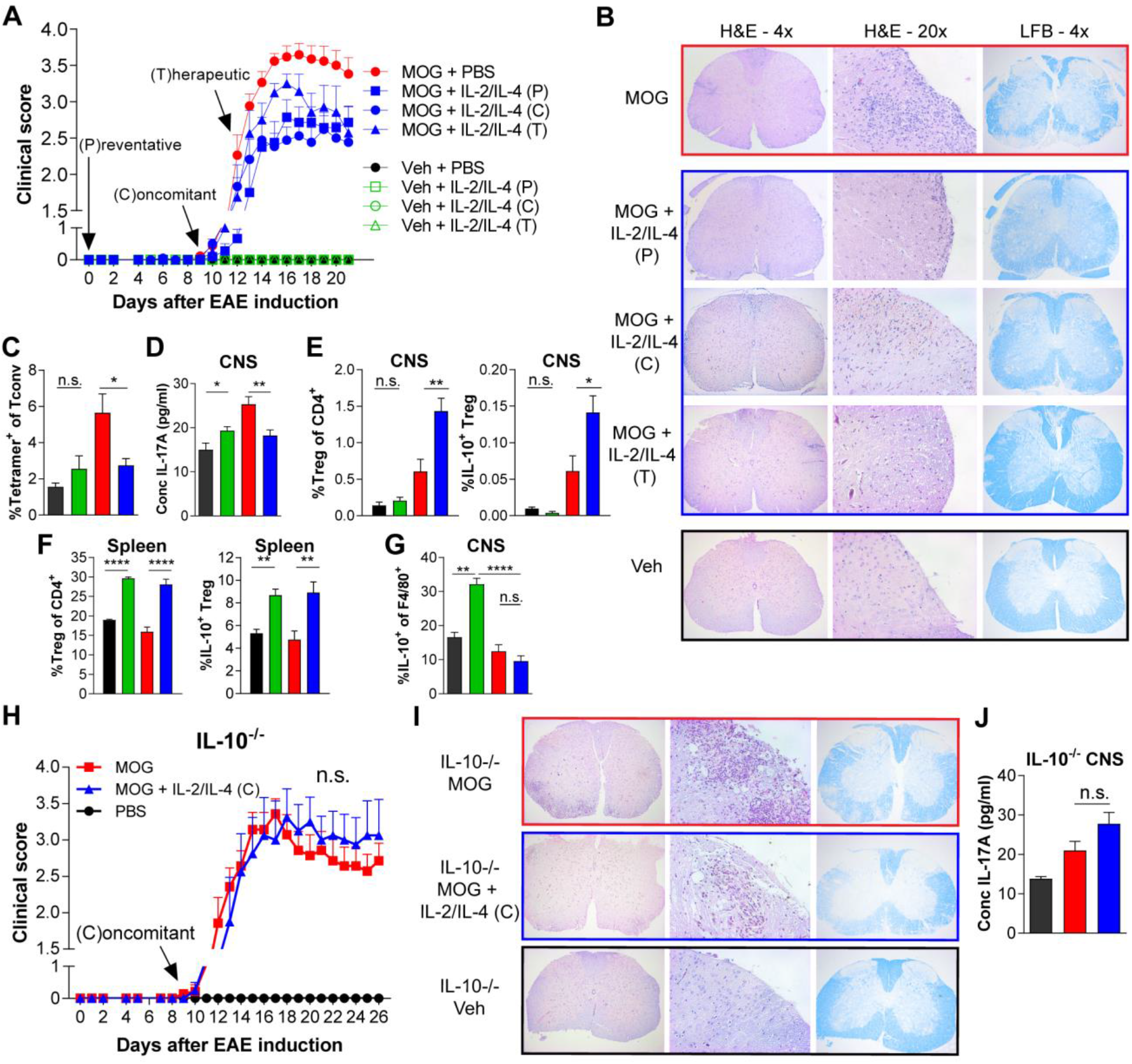
Combined IL-2 and IL-4 reduces the severity of EAE. (A-B) EAE clinical score and FFPE-processed spinal cords sectioned and stained with H&E and LFB from mice challenged with MOG peptide with and without combined cytokines (See also Figure 7 – figure supplement 1). Histological images represent a mouse of the mean disease score of each condition (See also Figure 7 – figure supplement 1C). N=10-12 for EAE mice; N=4 for vehicle control mice. (C) Analysis of MOG-loaded MHCII tetramer positive Tconv cells in the CNS as determined by flow cytometry. N=3 (D) Quantification of IL-17A secreted by CNS leukocytes isolated by Percoll gradient and incubated overnight in media, as quantified by ELISA. N=3 (E-F) Analysis of Treg abundance and IL-10-expression in Tregs isolated from the CNS or spleen of FoxP3^RFP^/IL-10^GFP^ mice in the EAE trials by flow cytometry. N=3 (G) Analysis of IL-10 expression of F4/80+ CNS macrophages from FoxP3^RFP^/IL-10^GFP^ mice in the EAE trials by flow cytometry. N=3 (H-I) Assessment of EAE severity in IL-10^-/-^ mice as quantified by daily clinical scoring and FFPE-processing of spinal cords stained with H&E and LFB. Images represent a mouse of the mean disease score of each condition. N=7-8 for EAE mice; N=4 for negative control mice for disease (J) Quantification of IL-17A secreted by CNS leukocytes of IL-10^-/-^ mice isolated by Percoll gradient and incubated overnight in media, as quantified by ELISA. N=3 For all panels, mean ± SEM are indicated. *p<0.05, **p<0.01, ****p<0.0001

To better understand the underlying mechanism of EAE suppression, we analyzed Tregs within the central nervous system (CNS) and spleen. We found that the cytokines in combination increased the percentage of CNS and splenic IL-10^+^ Tregs in MOG-challenged animals (Figures 7E and 7F) in correlation with reduced disease. Interestingly, we did not observe increases in IL-10 among CNS-localized F4/80+ macrophages in MOG-challenged mice with cytokine therapy (Figure 7G) despite the reported ability of IL-4 alone to induce IL-10 production in macrophages (*40*). Finally, we found that cytokine efficacy in EAE was lost in IL-10-deficient mice as measured by disease score (Figure 7H), cellular infiltration by H&E (Figure 7I) and IL-17A levels in the CNS (Figure 7J), thereby demonstrating the IL-10-dependency of combined IL-2 and IL-4 in suppressing EAE.

## Discussion

In this study, we discovered that the combinatorial signaling mediated by IL-2 and IL-4 in TCR-activated Tregs produced a response characterized by IL-10 expression synergy, selective proliferation of IL-10^+^ Tregs, differential STAT5 and STAT6 signaling, and distinct cell surface receptor phenotype compared to either cytokine in isolation. These factors cumulatively enhanced the ability of Tregs to suppress Tconv cell proliferation and cytokine secretion in co-culture while successfully attenuating inflammatory diseases of different underlying etiologies in preventative, concomitant, and therapeutic delivery regimens. Disease suppression was paired with increased Treg prevalence and IL-10 expression in Tregs, a reduction in MOG-specific effector T cells in EAE, and was abrogated in mice deficient in IL-10. Together, our data reveal that the combinatorial cytokine signaling in Tregs is integrated to generate a unique biological outcome that is divergent from the sum of each pathway in isolation and robustly enhances and maintains the suppressive ability of FoxP3+ Tregs *in vitro* and *in vivo.*

Previous studies have shown that two cytokines can cooperate by sharing the same STAT proteins to either enhance or interfere with the other to form a different outcome (*27, 41, 42*). While our study focuses on Tregs, others have reported optimal differentiation of Th2 cells with sequential IL-2 and then IL-4 stimulation, which upregulates IL-4Rα and is mediated by combined STAT5 and STAT6 phosphorylation (*43, 44*). Different from the sequential nature of these and other studies (*27*), our principle findings required simultaneous stimulation by both cytokines (Figure 1H-I). Furthermore, while FoxP3^-^ Tconv cells were reported to undergo a Th2-differentiating response to IL-4 and IL-2 (*43*), we found that this combination failed to induce IL-10 production in FoxP3^-^ Tconv cells (Figure 1A) while programming FoxP3^+^ Tregs towards an IL-10^+^ and suppressive phenotype. As a result, our data reveal that not only can the integration of multiple cytokine pathways diverge from the sum of the parts, but that the combinatorial nature can be different among varied cell types and lineages.

Interestingly, our data indicate that administration of IL-2 and IL-4 in mice *in vivo* suppressed the severity of not only IL-17-dependent EAE (*45*), but also several models of murine asthma in both Th1 and Th2-skewed backgrounds (*46*). This initially appears at odds with the notion that administration of IL-4 in asthmatic mice aggravates disease (*47*). However, we found that the combination of IL-2 and IL-4 develops Tregs with highly effective suppression of IL-4 production in Tconv cells *in vitro* (Figure 3). With the loss of IL-4-driven STAT6 phosphorylation in Tregs stimulated both cytokines, we have demonstrated that the IL-4 signal leading to STAT6 activation is dramatically inhibited upon simultaneous IL-2 exposure in Tregs. This reinforces the notion that the functional outcome of a cytokine is highly context dependent and may not reflect the outcome of the cytokine in isolation, leading to pleiotropy in complex systems.

In the case of IL-2 and IL-4, the net functional result of the discovered pathway is a greatly enhanced population of immune inhibitory IL-10^+^ Tregs. The cellular activity was potent enough to not only reduce disease prophylactically, but also to reverse ongoing inflammation in both asthma and EAE models. It is therefore interesting to note that the use of both IL-2 and IL-4 appears to be a characteristic of some beneficial microbiota antigen responses. For example, *Bacteroides fragilis,* which is native to a healthy human gut (*48, 49*), is known to regulate the peripheral immune system (*37, 38, 50, 51*) and even benefit the gut-brain axis (*52, 53*) through T cell recognition of its capsular polysaccharide PSA (*54*). It is now clear that PSA-specific effector T cells require communication with Tregs, and that immune suppression and inflammatory disease reversal depends upon both IL-4 (*29, 39*) and IL-2 (*55*), thereby providing a biological context in which the use of cytokines in combination has evolved to direct a healthy immune tone.

From a translational perspective, cytokine therapies have been limited (*56*) in part due to the short half-life in circulation (*57, 58*). As such, our findings suggest that IL-2 and IL-4-induced combinatorial signaling may be best harnessed for autologous Treg transfer therapies, which itself has been limited by poor expansion and survival of Tregs *ex vivo* (*59*). Here, Tregs that underwent stimulation by both IL-2 and IL-4 together not only retained their activated and IL-10-producing state, but also exponentially expanded and survived for almost two weeks in culture with 3 days of cytokine stimulation and 9 days of co-culture with Tconv cells (Figure 3J). Our data further suggests that the IL-2 and IL-4 combination reprograms Tregs with a particular receptor expression and STAT signaling pattern, potentially allowing for the cells to continue suppressing for extended periods through enhanced sensitivity to both cytokines.

In conclusion, we identified a novel mechanism of cell signaling integration controlling the transcriptional and proliferative response of regulatory T cells. While this is demonstrated in T cells using only two cytokines, these findings suggest that combinatorial signaling with other cytokines and cell types is likely. Moreover, while these differences in response may underlie cytokine pleiotropy, they also provide an intriguing guide for new therapeutic applications of combinatorial cytokines in the clinical setting ranging from *ex vivo* support of autologous cell transfers to immune activation in cancer and inhibition for autoimmunity.

## Materials and Methods

### Mice

C57BL/6J (Stock #000664), IL-10^eGFP^ (B6.129S6-Il10^tm1F/v^/J, Stock #008379), FoxP3^RFP^ (C57BL/6-FoxP3^tm1Flv^/J, Stock #008374), and IL-10-null (B6.129P2-//10^tm1Cgn^/J, Stock #002251) mice, all on the C57BL/6 background, as well as FoxP3^eGFP^ mice on the BALB/c background (Foxp3^tm2Tch^ Stock #006769) were purchased from the Jackson Laboratory (Bar Harbor, ME). IL-10^eGFP^ and FoxP3^RFP^ mice were crossed to make IL-10^eGFP^ x FoxP3^RFP^ dual-reporters in our facility. Mice were housed in a 12 hr light/dark cycle specific pathogen free facility and fed standard chow (Purina 5010) *ad libitum.* Enrichment and privacy were provided in mating cages by ‘breeding huts’ (Bio-Serv S3352-400). Mouse studies and all animal housing at Case Western Reserve University were approved by and performed according to the guidelines established by the Institutional Animal Care and Use Committee of CWRU.

### Primary splenic T cells

Primary splenocytes were isolated from freshly harvested mouse spleens, and reduced to a single cell suspension by passing them through a sterile 100μM nylon mesh cell strainer (Fisher Scientific, Hampton, NH). The single cell suspensions were labeled with anti-mouse CD4 magnetic microbeads (Miltenyi Biotec, San Diego, CA), and separated with an AutoMACS Pro Separator (Miltenyi Biotec) per manufacturer’s instructions.

### Cell culture

After flow sorting, Tregs or Tconv cells were cultured in 96-well round bottom plates (Corning, Corning, NY) at 50,000 cells per well in advanced RPMI (Gibco/Fisher Scientific, Waltham, MA) supplemented with 5% Australian-produced heat-inactivated fetal bovine serum, 55μM β-mercaptoethanol, 100U/mL and 100μg/mL Penicillin/Streptomycin, and 0.2mM L-glutamine (Gibco/Fisher Scientific, Waltham, MA) at 5% CO2, 37°C. For activating conditions, wells were coated with αCDЗε (eBioscience, San Diego, CA) at 2.5μg/mL in PBS then incubated at 4°C overnight followed by two washes with PBS before receiving cells. As indicated, cultures were supplemented with recombinant IL-2 (R&D Systems, Minneapolis, MN), IL-4 (R&D Systems), and IL-13 (Invitrogen, Carlsbad, CA), with equimolar concentration being 728 pM. For experiments involving STAT5 inhibitor, STAT5i (Stemcell, Vancouver, BC, CAN) resuspended in DMSO was added directly to culture at the designated amounts. Due to the known toxicity of DMSO, controls with DMSO only were also included, with % inhibition calculated compared to the DMSO controls.

### *In vitro* Tconv suppression

Tregs were pre-stimulated with αCDЗε and all combinations of cytokine for 3 or 7 days, washed, then mono-cultured or introduced into co-culture with 50,000 freshly isolated and CellTrace-stained Tconv cells at a 1:2, 1:4, 1:8, 1:16, 1:32, and 1:64 dilution (*60*). To assess the suppressive ability of a Treg on a per cell basis, Tregs were counted after stimulation and before placing in co-culture. To determine the suppressive ability of Tregs on a population level, Tregs were counted prior to stimulation so that the number of cells introduced into co-culture with Tconv reflected the proliferation induced by the cytokine conditions. CellTrace signal in Tconv cells was detected by flow cytometry after 3 days of co-culture, and secretion of cytokines were quantified by ELISA.

### EAE model of multiple sclerosis

Age-matched female mice between 11-23 weeks underwent EAE induction using the MOG_35-55_/CFA Emulsion and pertussis toxin (PTX) kit from Hooke Laboratories (Lawrence, MA) according to manufacturer’s instructions. Mice were immunized with 200 μg of MOG_35-55_/CFA emulsion s.c. in 2 locations in the back on day 0 and were administered 100 ng of PTX i.p. on days 0 and 1 of the trial. Negative controls for disease received a CFA emulsion and PTX according to the same schedule. Mice were weighed and scored daily according to Hooke Laboratory’s EAE scoring guide (https://hookelabs.com/protocols/eaeAI_C57BL6.html). Treated mice received cytokine cocktail comprising of equimolar IL-2 (20 ng) and IL-4 (16.24 ng) i.v. in a volume of 100 μl of sterile PBS every other day. Untreated control mice received 100 μl of sterile PBS i.v. at the same time intervals.

### HDM and TAC models of pulmonary inflammation

Age and sex-matched mice between 8-23 weeks were challenged with house dust mite antigen (HDM, *D. Farinae,* GREER, Lenoir, NC) by i.n. delivery of 20 μg HDM/dose in PBS according to the designated challenge schedule (Figures 5A, 5C, 6A) and sacrificed accordingly (*61*). IL-2 and IL-4 at the designated doses and time intervals were delivered i.n. in a mixed cocktail according after animals were anesthetized with 3% isoflurane (Baxter, Deerfield, IL) with an anesthesia system (VetEquip, Livermore, CA). In the chronic triple antigen combination model, mice were sensitized intraperitoneally with 20 μg Ovalbumin (OVA, Albumin from chicken egg white, Millipore Sigma, Darmstadt, Germany), 2.5 μg cockroach antigen (CRA, American, *Periplaneta Americana,* GREER, Lenoir, NC) and 2.5 μg of HDM in PBS. Immediately following sensitization, mice were challenged intranasally with HDM, CRA, or OVA in a rotating schedule for 7 weeks (*62*). For Flexivent (SCIREQ, Montreal, Quebec) studies, mice were anesthetized with a cocktail of Ketamine/Xylazine/Acepromazine, the trachea was exposed and catheterized, methacholine was administered via nebulization in increasing doses. Airway resistance was measured on a Flexivent respirator over time. After mice were euthanized, bronchial alveolar lavage fluid (BALf) was recovered in 3 lavages of 1 ml each, and lung tissues underwent FFPE processing or were reduced to a single cell suspension for flow cytometric analysis. BALf cell differentials were acquired by a HemaVet 950 Hematology Analyzer.

### Flow cytometry and cell sorting

For splenic Treg cell sorting, magnetic bead-mediated positively selected CD4+ cells were sorted by endogenously expressed FoxP3^RFP^ and/or IL-10^GFP^. For IL-10ko mice, CD4+ cells were stained and sorted with CD25-BV421 (BD Biosciences, San Jose, CA). Cells were washed in MACS buffer (Miltenyi Biotec) before sterile cell sorting using a FACSAria-SORP (BD Biosciences). For flow cytometric analysis of tissue compartments, cells were passed through a sterile 70μM nylon mesh cell strainer (Fisher Scientific, Hampton, NH) for homogenization into a single cell suspension. Blood and spleen were RBC-lysed (BD, Franklin Lakes, NJ), and brain and spinal underwent a 30%/70% Percoll gradient centrifugation (GE, Boston, MA). For intracellular staining, cells were fixed and permeabilized with the FoxP3/transcription factor staining buffer set (eBioscience). Commercial antibodies used included: CD4-PE (eBioscience), CD4 – AF647 (Biolegend), F4/80 – BV711 (BD), CD25-BV421 (BD), CD25-PE (eBioscience), CD25 – PE-Cy7 (BD), CD124 – BB700 (BD), CD122 – APC (BD), CD132 – BV421 (BD), CD213a1 – Biotin (MyBioSource, San Diego, CA), Biotin – BV-605 (BD), STAT3 – AF647 (BD), STAT5 – AF647 (BD), STAT6 – AF647 (BD) and mTOR – AF647 (BD). Additional flow cytometry reagents purchased commercially include: Sytox Red (Invitrogen), CellTrace Violet (Invitrogen), and Fixable Viability Stain 510 (BD). Tetramers obtained from the NIH Tetramer Core include MOG_35-55_ Tetramer – BV421, I-A(b) Tetramer – BV421 (Atlanta, GA). Cells were acquired on an Attune NxT cytometer (Invitrogen). Cytometer usage was supported by the Cytometry & Imaging Microscopy Core Facility of the Case Comprehensive Cancer Center. Analysis of all FACS data was performed using FlowJo v10 (Tree Star, Inc., Ashland, OR). Analysis of proliferation was done in FlowJo v10 using the Proliferation Modeling platform.

### Cell Proliferation

Cell proliferation was quantified using CellTrace Violet staining coupled with analysis of dilution using flow cytometry. Division Indices were calculated mathematically using the Proliferation Modeling platform in Flowjo by dividing the total number of cell divisions by the total number of cells in the starting culture:, where =generation. Proliferation Indices were calculated by dividing the total number of divisions by the number of cells that underwent division:.

### ELISA and Luminex

Cytokine concentrations were analyzed by sandwich ELISA performed according to manufacturer’s instructions (BioLegend). The assay was modified to utilize europium-conjugated streptavidin (Perkin-Elmer, San Jose, CA). Signal was detected with a Victor V3 plate reader (Perkin Elmer). For Luminex assays, media were snap frozen in liquid nitrogen and sent to Eve Technologies (Calgary, AB) for mouse 31-plex and TGFβ 3-plex analysis.

### Histology and microscopy

Harvested tissues were fixed in 10% formalin (VWR, Radnor, PA) for 24 hours and sent to AML Laboratories (Jacksonville, FL) for paraffin embedding. Embedded lung tissues were sectioned and stained with H&E, Periodic Acid Schiff (PAS), or trichrome by AML and spinal cord tissues were sectioned and stained with Hematoxyling and Eosin (H&E) (Ricca Chemical, Arlington, TX and Sigma Aldrich, St. Louis, MO) or Luxol Fast Blue (LFB) (Sigma Aldrich) in our lab. Images were acquired with a Leica DM IL LED (Wetzlar, Germany).

### Power Analyses

Hooke Laboratories established that 10-12 mice are needed in MOG challenge groups using their strain-specific standardized reagents and procedures to account for the natural scatter of this pre-clinical disease model. For the asthma models, we used historical means between positive and negative control groups and their associated standard deviation to establish statistical power in animal model experimental design. The power was set to 0.80 with α = 0.05 in a 2-sided test at https://www.stat.ubc.ca/~rollin/stats/ssize/n2.html). We found that 6 mice were sufficient per group.

### Data Analyses

All data are represented by mean ± SEM. Pulmonary inflammation data sets include a minimum of four, but more commonly six animals per group per experiment. EAE trials include four animals in the negative disease control conditions and 10-12 animals in groups that receive MOG peptide. Data and statistical measurements were generated with Prism (Graphpad, San Digo, CA). For comparisons between two groups, Student’s t-test was used; comparisons between multiple groups utilized analysis of variance. The Principle Component analysis was generated in RStudio using factoextra and ggplot2. Mean ± SEM are indicated. *p<0.05, **p<0.01, ***p<0.001, ****p<0.0001.

## Supplementary Materials

**Figure 1 – figure supplement 1.** Cell sorting gating strategy and post-sort purity

**Figure 1 – figure supplement 2.** IL-10 is the only synergistic and robust analyte produced by Tregs following combinatorial cytokine stimulation

**Figure 1 – figure supplement 3.** Dynamics of combinatorial cytokine stimulation of Tregs over time

**Figure 4 – figure supplement 1.** The IL-10 and proliferative response following combined IL-2 and IL-4 stimulation is dependent on STAT5 signaling

**Figure 6 – figure supplement 1.** Administration of IL-2 and IL-4 suppresses HDM-induced acute asthma and is dependent on IL-10

**Figure 6 – figure supplement 2.** Administration of IL-2 and IL-4 therapeutically suppresses TAC-induced chronic asthma

**Figure 7 – figure supplement 1.** Combined IL-2 and IL-4 suppressed the severity of EAE in Preventative, Concomitant, and Therapeutic dosing regimen

## Acknowledgments

We are immensely grateful for the help of Douglas M. Oswald, PhD, for valuable scientific input and guidance in R, Mark B. Jones, PhD for mentorship and project discussions early on, Kalob M. Reynero for technical assistance in tissue sectioning, Jill M. Cavanaugh for technical assistance and maintenance of the mouse colony, and Lori. S.C. Kreisman for laboratory support and manuscript editing. Moreover, we thank Sandra Siedlak and Xiongwei Zhu, PhD for expertise in histological sectioning and staining, and Alex Huang, MD, PhD for equipment for histological imaging. We thank the Cytometry & Microscopy Shared Resource of the Case Comprehensive Cancer Center (P30CA043703) for equipment and assistance with flow cytometry-based experiments. This work was made possible by grants from: The National Institutes of Health (R01-GM115234), and the Hartwell Foundation to BAC, and the National Institutes of Health (T32-AI089474) to JYZ and CAA.

## Author contributions

Conceptualization, J.Y.Z. and B.A.C.; Methodology, J.Y.Z. and B.A.C.; Investigation, J.Y.Z. and C.A.A.; Formal Analysis, J.Y.Z. and B.A.C.; Writing – Original Draft, J.Y.Z.; Writing – Review & Editing, J.Y.Z. and B.A.C.; Funding Acquisition, B.A.C.; Resources, B.A.C.; Supervision, B.A.C.

## Abbreviations

IL-2: interleukin-2
IL-4: interleukin-4
IL-10: Interleukin-10
IL-17: interleukin-17
IL-2RγC: IL-2 receptor common gamma chain, CD132
IL-2Rα: IL-2 receptor alpha chain, CD25
IL-2Rβ: IL-2 receptor beta chain, CD122
IL-4Rα: IL-4 receptor alpha chain, CD124
IL-13Rα1: IL-13 receptor subunit alpha 1, CD213α1
Treg: regulatory T cell
Tconv: conventional T cell
EAE: experimental autoimmune encephalomyelitis
FoxP3: forkhead box P3
RFP: red fluorescent protein
GFP: green fluorescent protein
TCR: T cell receptor
PCA: principle component analysis
HDM: house dust mite
CRA: cockroach antigen
OVA: ovalbumin
TAC: triple antigen cocktail
BALf: bronchial alveolar lavage fluid
H&E: hematoxylin and eosin
PAS: periodic acid Schiff
LFB: luxol fast blue
MOG: myelin oligodendrocyte glycoprotein
CFA: complete Freund’s adjuvant
PTX: pertussis toxin
CNS: central nervous system
PSA: polysaccharide A

## Supplementary Materials

**Figure 1, figure supplement 1.**
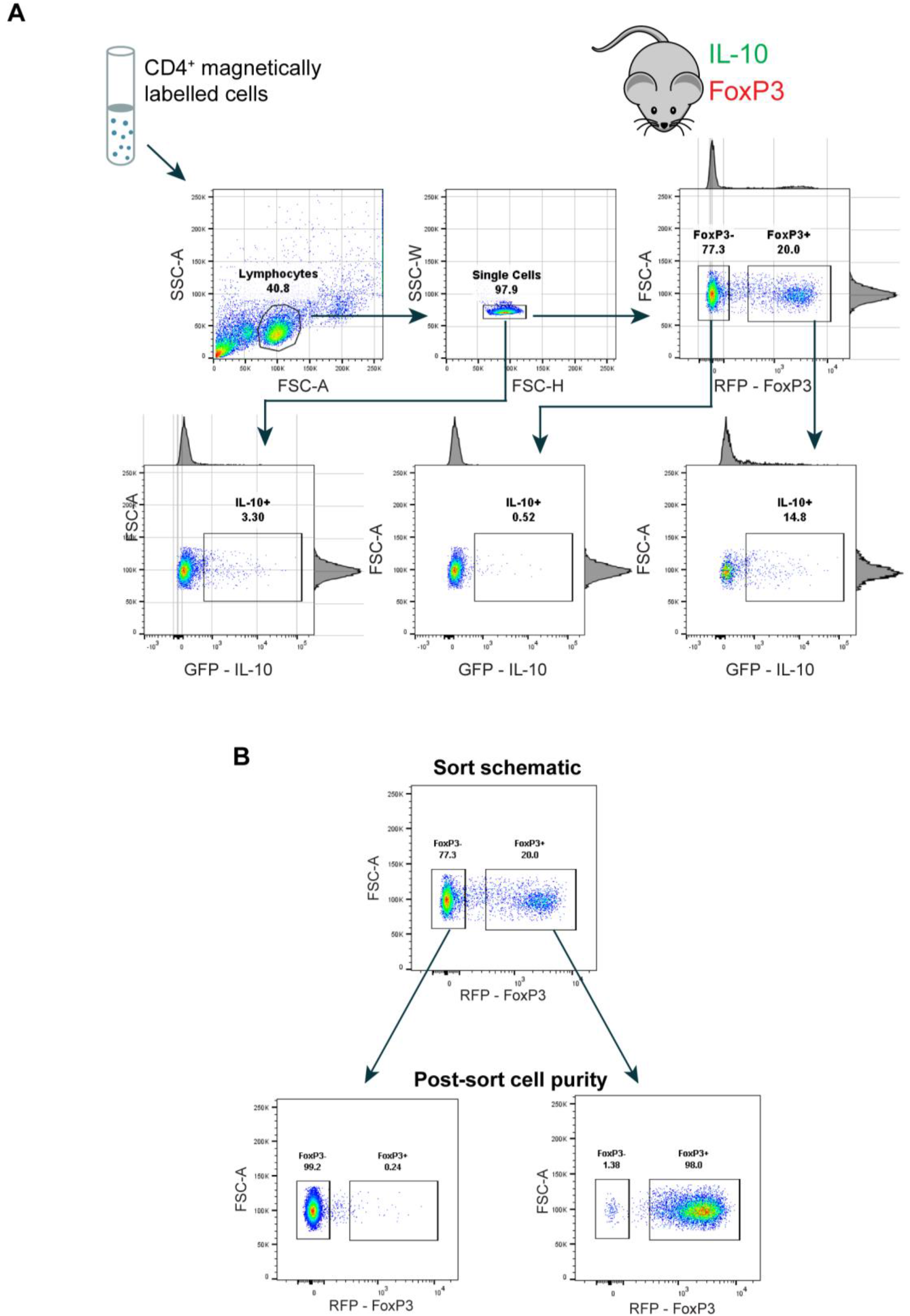
Cell sorting gating strategy and post-sort purity. (A) Gating strategy for flow cytometry sorting and analyzing IL-10^+/-^ Tregs from FoxP3^RFP^/IL-10^GFP^ dualreporter mice. Freshly harvested spleens underwent CD4^+^ magnetic bead selection prior to FACS. Related to Figure 1. (B) Sorted cells were immediately analyzed by flow cytometry to determine purity. Related to Figure 1.

**Figure 1, figure supplement 2.**
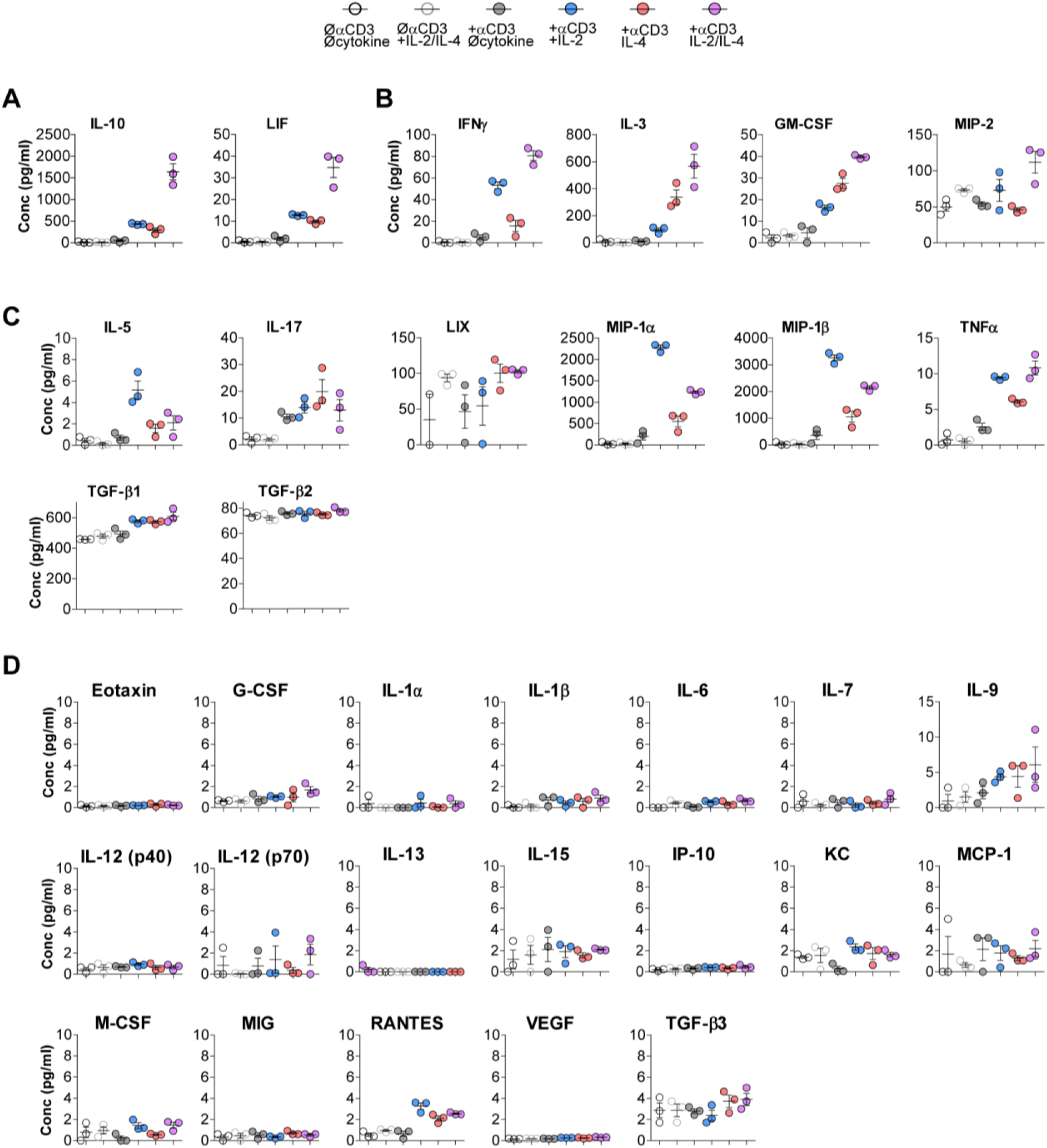
IL-10 is the only synergistic and robust analyte produced by Tregs following combinatorial cytokine stimulation. 31-plex Luminex analysis of cytokines secreted into the media following the designated stimulation conditions grouped in cytokines that (A) synergistically increase with combinatorial cytokine stimulation, (B) are summative, (C) show no change compared to single cytokine stimulation, (D) are not highly expressed. Error bars indicate mean ± SEM. N=3. Related to Figure 1J.

**Figure 1, figure supplement 3.**
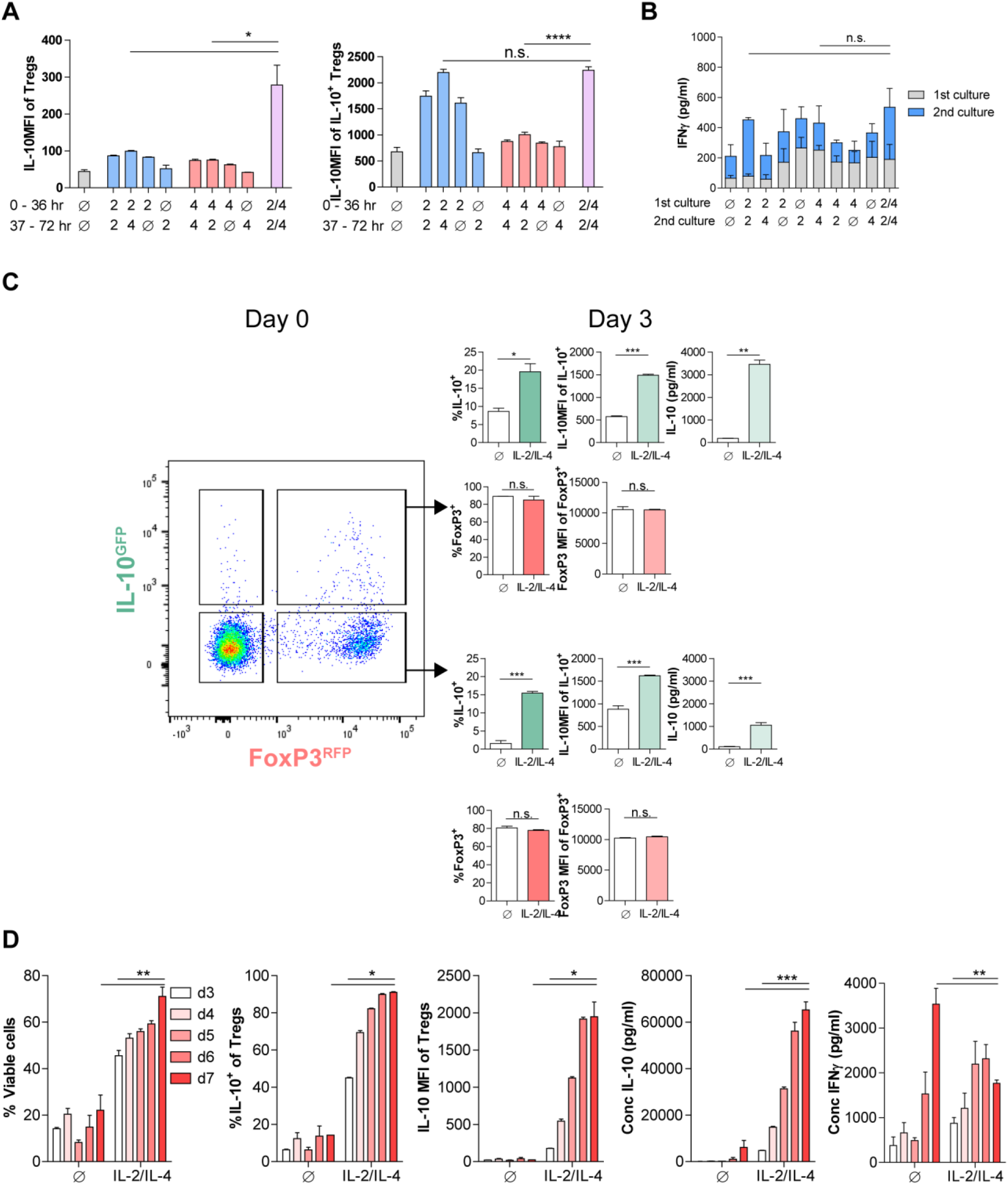
Dynamics of combinatorial cytokine stimulation of Tregs over time. (A) IL-10 expression of purified Tregs stimulated for 36 hours in culture with the first set of conditions, washed, and subsequently stimulated for another 36 hours in culture with the second set of conditions, as analyzed by flow cytometry. All samples received αCD3ε activation. N=3. Related to Figure 1H (B) Quantification of IFNγ secretion following the culture conditions as detailed above, as analyzed by ELISA. N=3. Related to Figure 1I. (C) IL-10 and FoxP3 expression of purified IL-10^+/-^ Tregs that underwent culture with αCDЗε and combined IL-2/IL-4for 3 days, as analyzed by flow cytometry. N=3. Related to Figure 1L. (D) Analysis of viability and IL-10 expression of purified Tregs cultured with αCDЗε and combined IL-2/IL-4 for 3-7 days, as analyzed by flow cytometry. IL-10 and IFNγ content in the culture media was analyzed by ELISA. For all panels, mean ± SEM are indicated. *p<0.05, **p<0.01, ***p<0.001, ****p<0.0001

**Figure 4, figure supplement 1.**
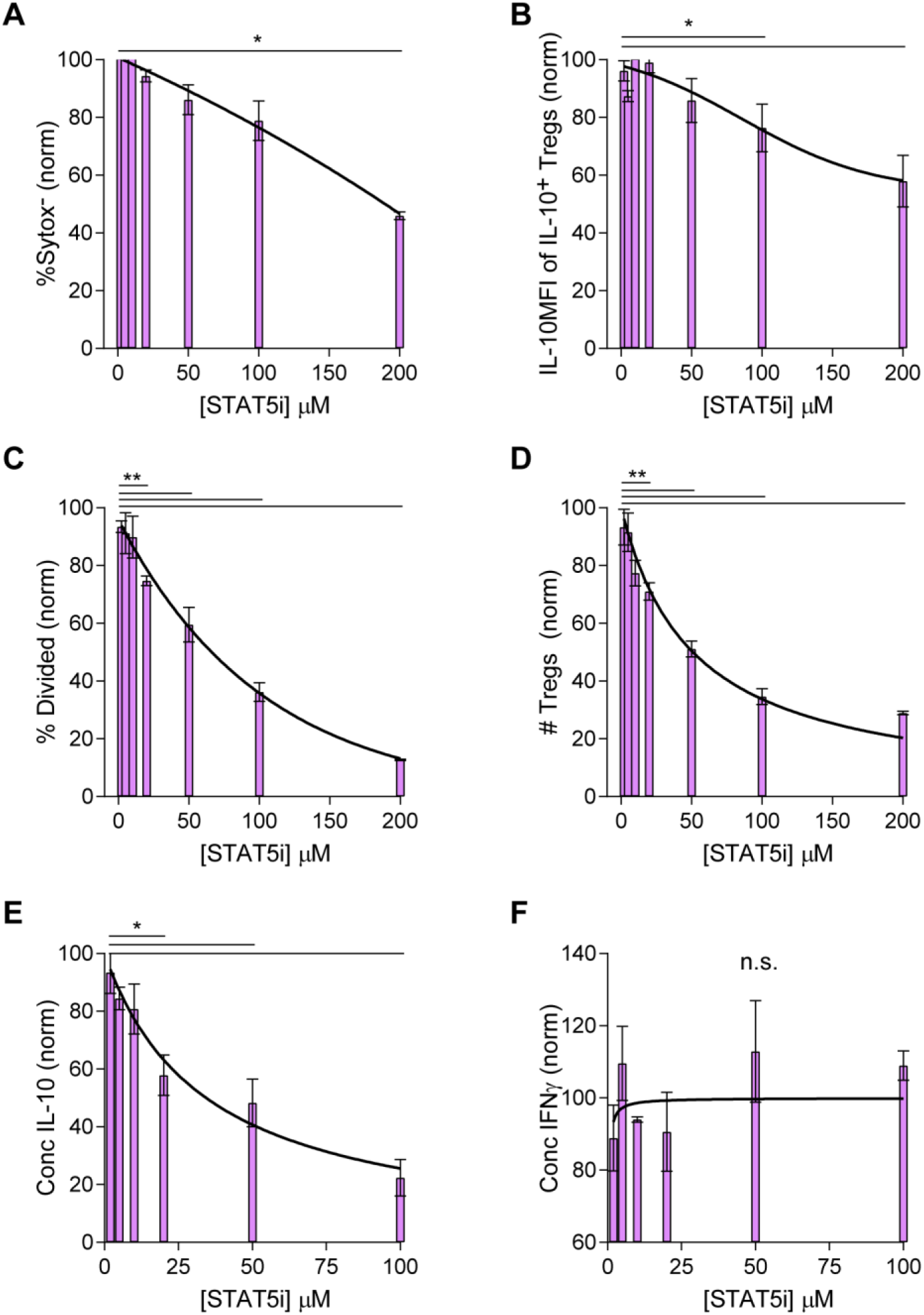
The IL-10 and proliferative response following combined IL-2 and IL-4 stimulation is dependent on STAT5 signaling. (A-D) Analysis of viability (Sytox Red), IL-10 expression, and proliferation of purified Tregs that were cultured with αCDЗε, combined IL-2 and IL-4, and STAT5i (resuspended in DMSO) in the indicated concentrations for 3 days, as analyzed by flow cytometry. The values are normalized to Tregs cultured with the same stimulation conditions but with DMSO only and no STAT5i. (E-F) Quantification of IL-10 and IFNγ secreted by the cells in the culture setup above as analyzed by ELISA. N=3. Related to Figure 4. For all panels, mean ± SEM are indicated. *p<0.05, **p<0.01

**Figure 6, figure supplement 6.**
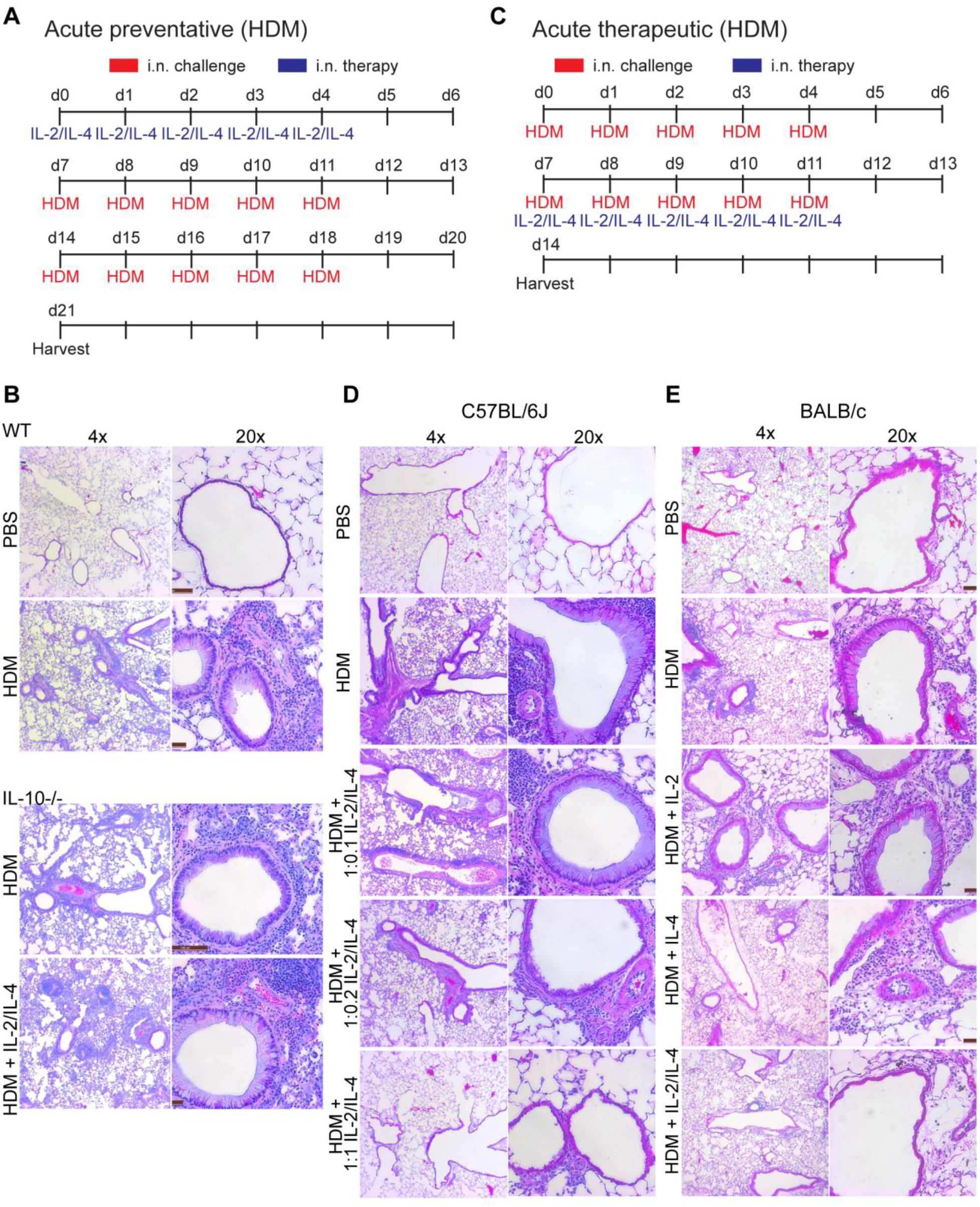
Administration of IL-2 and IL-4 suppresses HDM-induced acute asthma and is dependent on IL-10. (A) Challenge and treatment schedule for preventative HDM-induced acute asthma trial in Figures 6A and 6E. IL-2/IL-4 was delivered i.n. for 5 consecutive days (d0-5), followed by 2 weeks of HDM (or PBS) challenge through i.n. challenge (d7-11, d14-d18). (B) Representative H&E histology for Figure 6E. N=6 mice. (C) Challenge and treatment schedule for therapeutic HDM-induced acute asthma trial in Figure 6F. Mice were challenged with HDM (or PBS) for 2 weeks (d7-11, d14-18). IL-2/IL-4 was administered i.n. for 5 days starting on d7. (D) The severity of pulmonary disease was analyzed by FFPE tissues that underwent H&E staining and imaging at 4x and 20x. N=6 mice. Representative H&E histology for Figure 6F. (E) Pulmonary disease was analyzed by FFPE tissues that underwent sectioning, H&E staining, and imaging at 4x and 20x. N=6 mice. Representative H&E histology for Figure 6G.

**Figure 6, figure supplement 2.**
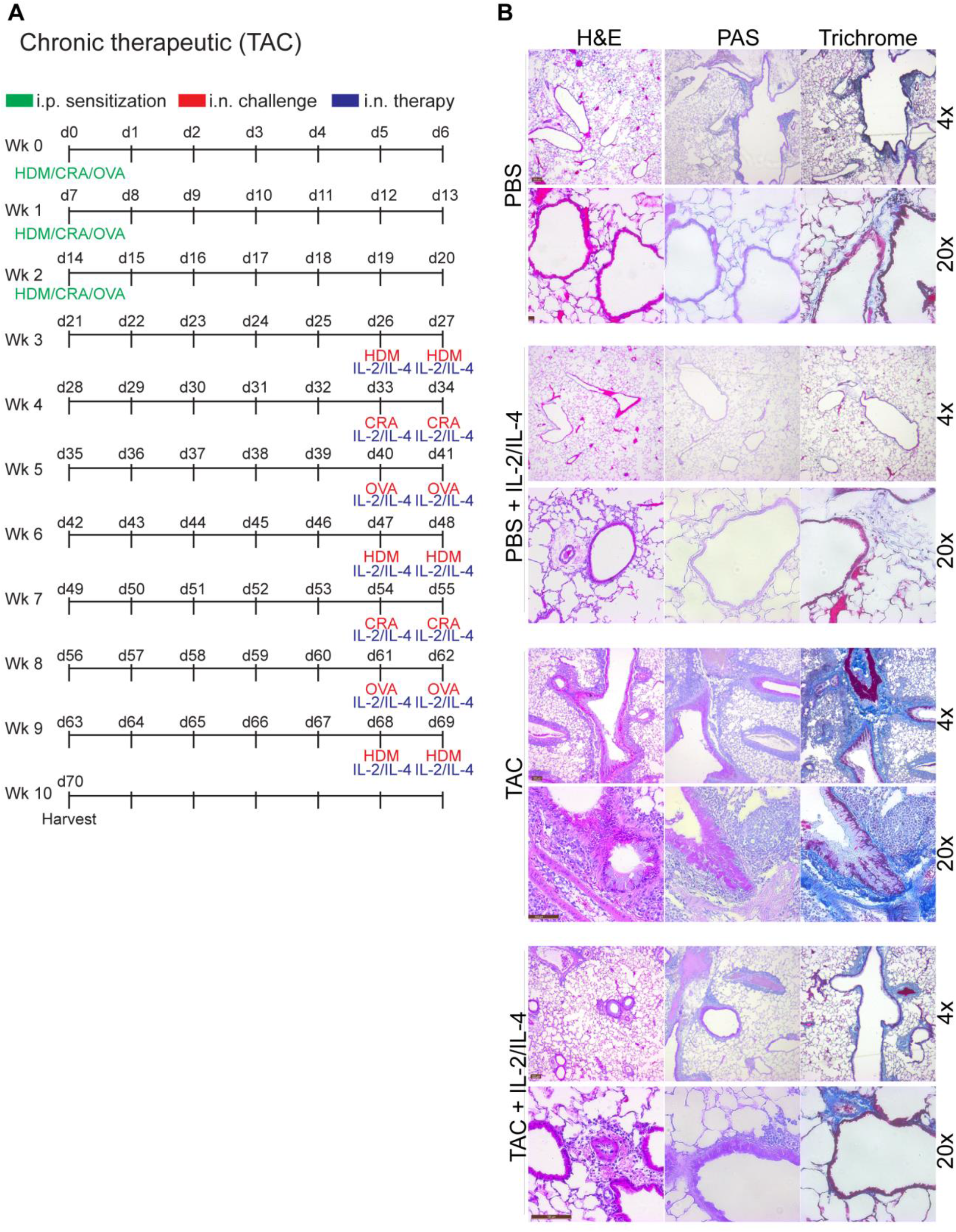
Administration of IL-2 and IL-4 therapeutically suppresses TAC-induced chronic asthma. (A) Challenge and treatment schedule for therapeutic TAC-induced chronic asthma trial in Figure 6H. Mice were sensitized to the triple antigens once weekly for the first three weeks. Afterwards, all challenges and treatments were delivered i.n. Mice received HDM, CRA, and OVA in a rotating weekly schedule, and also received IL-2 and IL-4 in combination following the challenge. (B) Pulmonary disease severity was analyzed by FFPE tissues that underwent H&E, PAS, Trichrome staining and imaged at 4x and 20x. N≤5. Representative H&E histology for Figure 6H.

**Figure 7, figure supplement 1.**
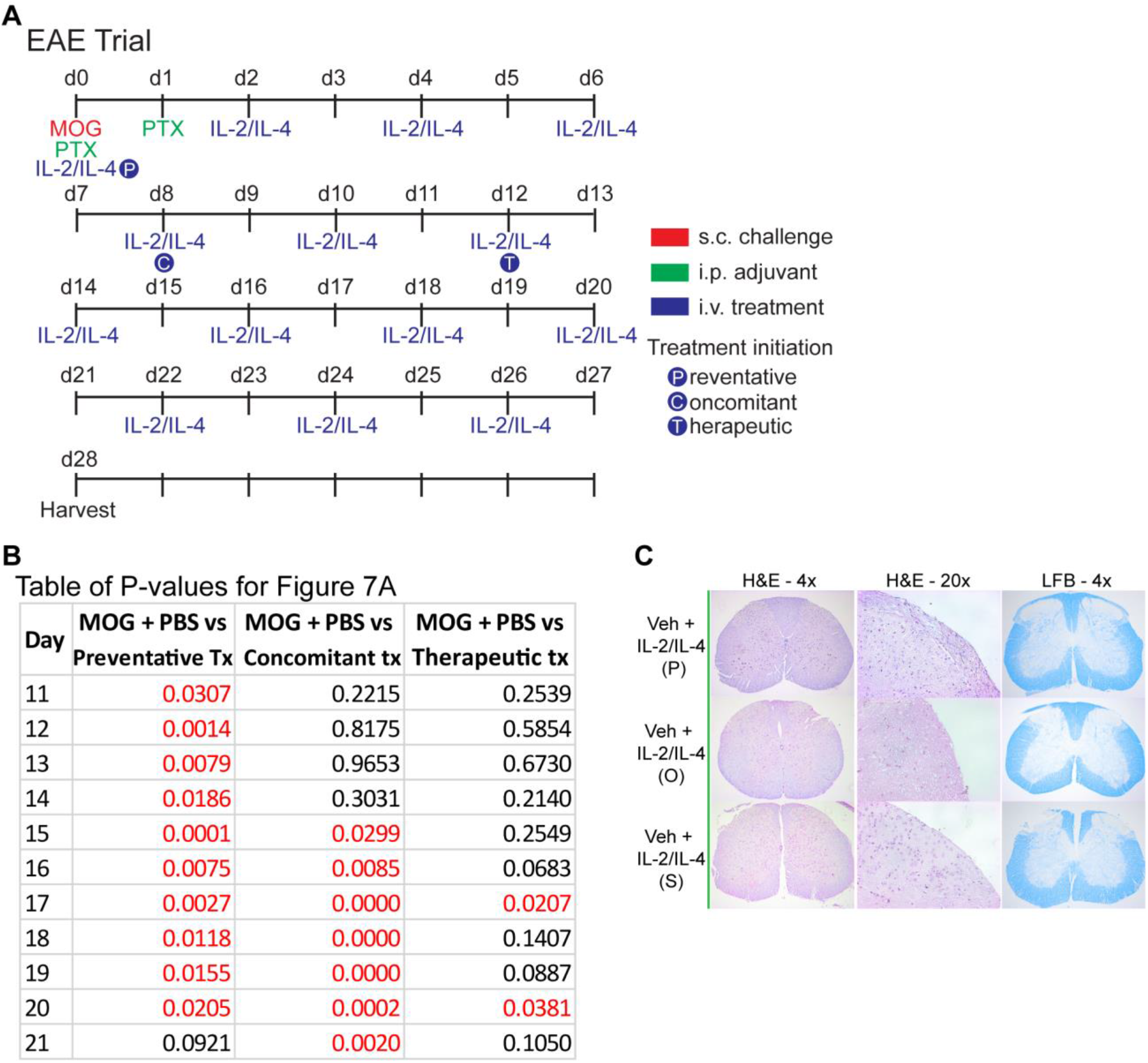
Combined IL-2 and IL-4 suppressed the severity of EAE in Preventative, Concomitant, and Therapeutic dosing regiment. (A) EAE was induced in mice with s.c. injections of MOG emulsified in Complete Freund’s Adjuvant (CFA) and d0. Pertussis toxin (PTX) was injected i.p. on days 0 and 1. Negative control mice received CFA only with PTX. The combinatorial cytokines were delivered i.v., with the Preventative regimen beginning on d0, Concomitant regimen on d8, and Therapeutic regimen on d12. Mice that did not receive the combinatorial cytokines received i.v. PBS as the vehicle control. Related to Figure 7. (B) Table of P values for Figure 7A. Red font represents P≤0.05. Black font represents P>0.05. N=10-12 for EAE mice, N=4 for vehicle control mice. Related to Figure 7A. (C) Analysis of disease in the mice that received vehicle controls through FFPE-processed spinal cords undergoing H&E or LFB staining. Images represent a mouse of the mean disease score of each condition. N=4 for vehicle control mice. Related to Figure 7B.

